# Joint tissue explant model using psoriatic arthritis synovial fluid as a tool to capture patient-specific responses to treatments

**DOI:** 10.64898/2026.07.05.736607

**Authors:** Atoosa Ziyaeyan, Mozhgan Rasti, Rajiv Gandhi, Katerina Oikonomopoulou, Vinod Chandran, Sowmya Viswanathan

## Abstract

**Objective:** We developed a patient- and joint-specific explant co-culture system to model active psoriatic arthritis (PsA) and capture donor-specific tissue responses to therapeutic interventions.

**Methods:** Based on convergent joint pathology between end-stage osteoarthritis (OA) and PsA, OA cartilage-bone and synovium tissues from arthroplasty patients were exposed to synovial fluid (SF) obtained from PsA and OA patients. Histological outcomes (synovitis, proteoglycan distribution), curated gene expression, soluble mediators, and proteinase activity were assessed over 7-21-days. Model responses to dexamethasone (DEX) and the anti-tumor necrosis factor antibody adalimumab (ADA) were evaluated.

**Results:** PsA SF induced distinct inflammatory and tissue remodeling responses compared to OA SF and control conditions, including altered cartilage proteoglycan distribution, increased synovitis, and tissue-specific transcriptional changes. Multivariate analyses identified distinct osteochondral and synovial transcriptional responses to PsA SF, characterized by reduced osteochondral *COL2A1* expression and increased synovial expression of inflammatory and matrix-remodeling genes, including *MMP1* and *CXCL8*. DEX and ADA elicited donor-specific responses across histological, transcriptional, and protein readouts. Among multivariable model outputs, histologic synovitis scores emerged as the most clinically aligned parameter, demonstrating associations with baseline PsA donor disease activity, active joint counts, pain, high-sensitivity C-reactive protein (hsCRP), and radiographic scores. Synovitis score changes to DEX and ADA treatments also aligned with corresponding PsA SF donor clinical improvements to corticosteroid and TNF-modifying therapies.

**Conclusion:** This osteochondral-synovial explant co-culture model captured donor-specific inflammatory and treatment-responsive features of PsA SF-induced pathology, thereby providing a clinically relevant ex vivo platform for studying patient-specific therapeutic responses in PsA.

## Introduction

Psoriatic arthritis (PsA) is a chronic inflammatory disease affecting musculoskeletal tissues and skin in about 25% of people with psoriasis (1,2). Despite the availability of targeted therapies, persistent disease activity remains a problem, with only a third of patients on targeted therapy and a fifth on methotrexate achieving minimal disease activity (3). Only 40% of PsA patients treated with biologic agents against tumor necrosis factor (TNF) or interleukins (IL)-12,-23,-17, or with small molecule inhibitors targeting Janus kinases or phosphodiesterase 4 achieve an American College of Rheumatology 20% (ACR20) response (4). Therefore, the management of therapy-resistant disease remains a challenge, stressing the need for further efforts in personalized medicine approaches to customize patient-specific treatments.

There are a limited number of *in vitro* PsA models, including those that utilize primary CD4^+^ T helper cells, peripheral blood mononuclear cells or synovial fluid synoviocytes from PsA patients; they have been useful to study Th17 and osteoclastogenesis in PsA, but do not fully capture pathophysiological mechanisms in the joint tissues (5,6). There are several useful *in vivo* models for studying various aspects of PsA pathology (7–11), but these have limited utility in effectively capturing the heterogeneity in PsA patient disease and responses to treatments. Developing *ex vivo* models of PsA remains an unmet need.

Joint disease in PsA overlaps with osteoarthritis (12,13), including patterns of joint involvement, initiation by (micro)trauma, inflammation, monocyte infiltration, cartilage degradation, new bone formation, involvement of innate and adaptive immune cells and multiple signaling pathways. PsA joint biopsies are not routinely obtained as it is an invasive procedure with risk of infection (14). Therefore, as an alternative, we used OA explanted bone, cartilage and synovium joint tissues with mismatched PsA donor SF to recapitulate end-stage PsA joint disease; OA explant tissues are routinely obtained during total knee arthroplasty (TKA). However, OA is considered less inflammatory, and OA samples are commonly used as controls for PsA studies. To mitigate these limitations, we used cell-free synovial fluid (SF) from PsA patients, which contains soluble effectors from monocytes, macrophages, T cells, and neutrophils, the primary drivers of joint inflammation (12,15). SF is an ultrafiltrate of blood admixed with components from joint tissues and other factors essential for maintaining joint homeostasis (16); changes to molecular and cellular components of the SF recapitulate inflammatory and damage-related pathogenic processes in the joint as previously shown (17).

The translational utility of this model was further supported by its responsiveness to clinically relevant anti-inflammatory therapies, including dexamethasone (DEX) and the anti-TNF antibody adalimumab (ADA). As such, this platform may serve as a patient-specific *ex vivo* tool to evaluate therapeutic responses and support personalized treatment strategies in PsA.

## Methods

Detailed experimental procedures, treatment conditions, reagent information, and expanded statistical analyses are provided in the Supplementary Methods.

### Tissue Collection and Preparation

Cartilage, synovium, and bone were obtained from 14 end-stage KOA patients (Kellgren-Lawrence grades 3-4) undergoing TKA, with informed consent (REB 14-7483). Seven OA SF samples were collected and pooled (REB 14-7483). Ten PsA SF were also collected (REB 21-5591). A single PsA patient undergoing TKA provided matched tissue and SF (REB 21-5591). Donor demographics are provided (Suppl Table 1-2).

### Explant Processing and Co-culture Model

Cartilage-bone explants were generated using a 2.7 mm osteochondral grafting system (Smith & Nephew). Synovium explants were trimmed and cultured on 0.4 μm transwell inserts. Explants were maintained in co-culture following 48-hour acclimatization. and cultured for up to 21 days with medium replenishment twice weekly.

### Histology

Explant tissues were fixed, paraffin-embedded, sectioned (4 μm), and stained using Safranin O/Fast Green or hematoxylin and eosin (H&E). Cartilage proteoglycan distribution was quantified using image-based analysis in FIJI (ImageJ software, Bethesda, USA), and synovitis severity was assessed using the Krenn synovitis scoring system (18).

### RNA Isolation and Gene Expression Analysis

RNA isolation and qPCR were performed as previously described (13). Gene expression was normalized to housekeeping genes and expressed as -ΔΔCt using tissue-specific gene panels (Suppl Table 3).

### Soluble Mediator and Western Blot Analyses

IL-6, CCL2, and total matrix metalloproteinase (pan-MMP) activity were quantified using commercially available ELISA and fluorometric activity assays. Western blotting was performed to assess total and phosphorylated c-Jun N-Terminal Kinase (JNK) expression. Raw protein measurement data and original Western blot images are provided in Suppl Table 4 and Fig S1A-B, respectively.

### Statistical Analysis

Statistical analyses were performed using JMP Pro v17 (SAS Institute). Group comparisons were performed using parametric or non-parametric tests as appropriate based on normality checks. Correlations were assessed using Spearman’s rank correlation. Multivariate analyses included principal component analysis (PCA) and canonical discriminant analysis (DA). Data are presented as mean ± standard deviation (SD). A p-value < 0.05 was considered statistically significant, and false discovery rate (FDR) correction was applied where appropriate.

## Results

### OA Explant Co-culture model with PsA Synovial Fluid Induces Distinct Inflammatory and Degradative Signatures

A co-culture model using osteochondral and synovium explant tissues from end-stage OA patients undergoing TKA was supplemented with PsA SF and compared to control groups, including a baseline co-culture (negative control) and pro-inflammatory cytokine-stimulated positive controls (Fig 1A). Unsupervised hierarchical clustering revealed distinct separation of PsA vs. control groups after 7 days of co-culture (Fig 1B). While the model demonstrated robust and distinct osteochondral and synovium gene expression profiles with PsA SF stimulation through day 21, a 7-day time point was selected for practical reasons, as it required lower PsA SF volumes. Additionally, on days 7, 14 and 21, osteochondral and synovium gene expression profiles largely overlapped, based on unsupervised cluster analysis (Fig S2A-B). Multiple concentrations (%v/v in explant culture medium) of PsA SF were also tested, but in the absence of a specific dose response (Fig S3A-B), a feasible 30% (%v/v) was chosen.

**Figure 1.**
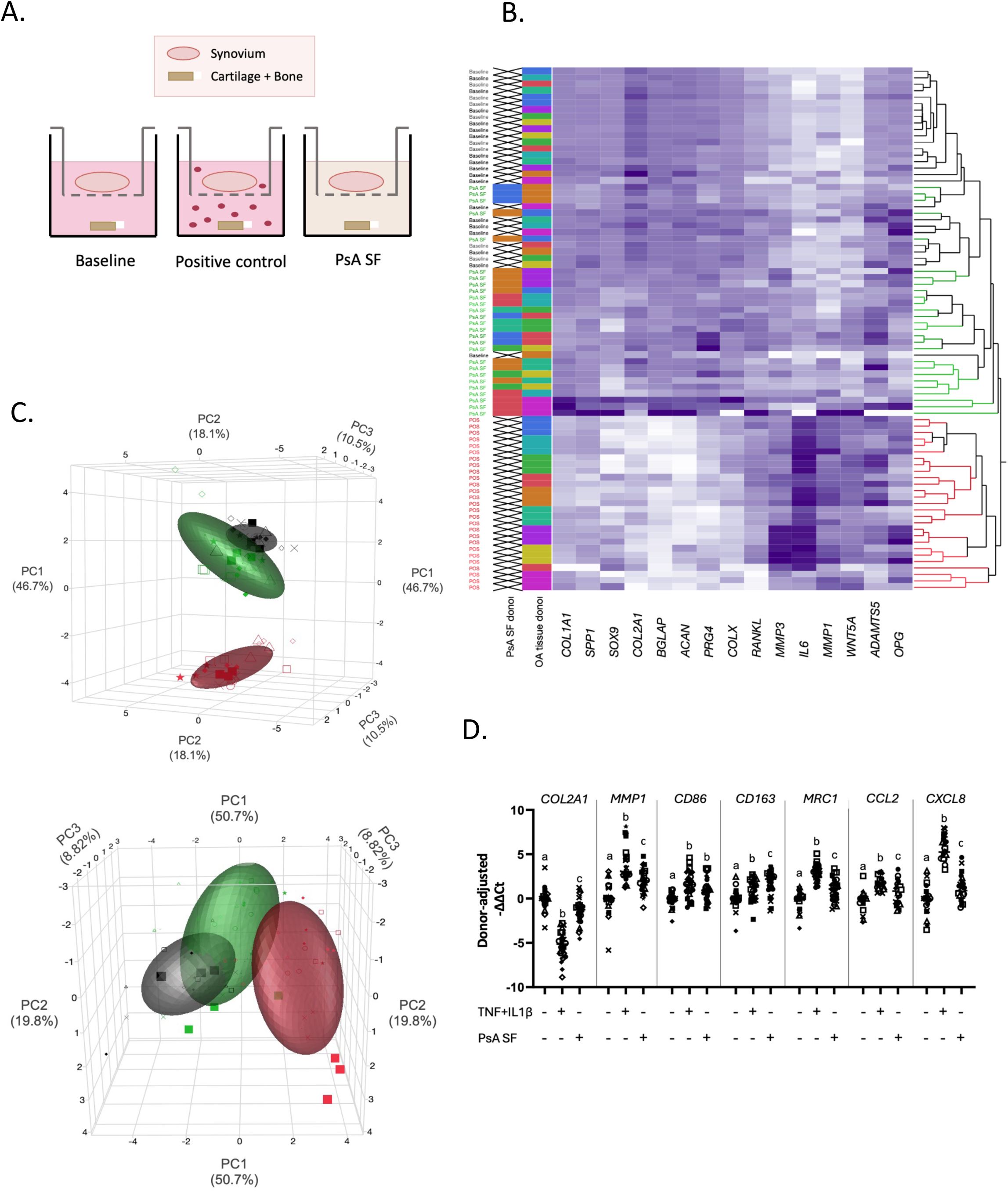
Osteochondral-synovial co-culture model demonstrates distinct transcriptional responses following PsA synovial fluid stimulation. **(A)** Schematic of the cartilage-bone (CB) and synovium (SYN) co-culture system with baseline, TNF + IL-1β positive control (POS), and psoriatic arthritis (PsA) synovial fluid (SF) supplemented conditions. **(B)** Heatmap of CB gene expression across experimental conditions, each performed in technical triplicate. Gene expression values were column-standardized prior to unsupervised hierarchical clustering using Ward’s linkage method. Side bar annotations indicate experimental condition, PsA SF donors (n = 5), and OA tissue donors (n = 10). **(C)** 3D principal component analysis (PCA) of CB (top) and SYN (bottom) gene expression profiles. Each point represents one co-culture sample of a single OA tissue donor exposed to the indicated experimental conditions. Ellipses represent group clustering distributions. Black, baseline; red, POS; and green, PsA conditions. Distinct symbols correspond to individual OA tissue donors (Suppl Table 1). Variance is indicated on the axes; PCA loading coefficients are provided (Suppl Table 5). **(D)** Relative gene expression (donor-adjusted -ΔΔCt) of selected CB and SYN markers across experimental conditions following statistical blocking for OA tissue donor effects, in triplicate. Values were normalized to respective baseline controls. Different letters indicate statistical significance (one-way ANOVA with Tukey’s post hoc testing, p < 0.05).

PCA of osteochondral and synovium gene expression confirmed unsupervised hierarchical analysis, demonstrating unique clusters of treatment groups within each panel (Fig 1C). Euclidean distances between group centroids showed that the PsA SF-treated tissues clustered separately from the basal (1.43-cartilage-bone; 2.03-synovium) and positive control (5.4-cartilage-bone; 3.11-synovium) co-cultures, suggesting a PsA SF-specific transcriptome in both tissues.

In the osteochondral gene dataset, PC1 (46.7% variance) captured an axis of structural tissue homeostasis (*BGLAP*, *ACAN*, *COL2A1*, *COL1A1*, and *PRG4*) vs. inflammatory and catabolic mediators (*IL6*, *MMP1*, and *MMP3*). PC2 (18.1%) and PC3 (10.5%) reflected secondary variations related to osteochondral remodeling and bone turnover (Suppl Table 5). In the synovium dataset, PC1 (50.7% variance) reflected broad inflammatory and macrophage activation signatures, with major contributions from *CD206*, *CXCL8*, *MMP1*, *IL1B*, and *CCL2* (Fig. 1C). PC2 (19.8%) captured macrophage polarization-related variations, while PC3 (8.82%) reflected an *IL-17*-associated inflammatory signature dominated by *IL17A* (Suppl Table 5). Notably, in the osteochondral tissue, PsA SF significantly decreased *COL2A1* expression compared to baseline control, and to a greater extent than the pro-inflammatory cytokine-supplemented positive control (Fig 1D). In the synovium, PsA SF upregulated *MRC1*, *CD163*, *CD86*, *CCL2*, *MMP1*, and *CXCL8,* albeit to similar or lesser extents than the pro-inflammatory cytokine positive control (Fig 1D). Compared to the untreated controls, the addition of PsA SF resulted in modest increases in IL-6 and CCL2 protein levels relative to baseline (Table 1). By contrast, pan-MMP activity level showed substantial donor variability and did not consistently parallel transcriptional changes.

**Table 1.** Fold change in IL-6, CCL2, and pan-MMP enzymatic activity levels in conditioned medium. Values are presented as mean ± SD fold change relative to baseline controls, with additional normalization to the corresponding SF-alone conditions. N=3 OA tissue exposed to n=2 PsA SF donors for DEX, and n=3 PsA SF donors for ADA treatments; n=1 PsA tissue exposed to a matched PsA SF donor, in triplicate. IL-6 = interleukin-6; CCL2 = C-C motif chemokine ligand 2; MMP = matrix metalloproteinase; OA = osteoarthritis; PsA = psoriatic arthritis; SF = synovial fluid; DEX = dexamethasone; ADA = adalimumab.

|  | **Condition** | **IL-6** | **CCL2** | **MMP activity** |
| --- | --- | --- | --- | --- |
| **Osteoarthritis**<br>**explant tissues** | Baseline | 1.00 | 1.00 | 1.00 |
|  | Pro-inflammatory<br>cytokine control | 21.06 ± 9.07 | 11.39 ± 5.11 | 21.99 ± 29.95 |
|  | PsA SF | 1.92 ± 0.59 | 1.55 ± 0.33 | -5.97 ± 9.77 |
|  | PsA SF +DEX | 1.40 ± 0.48 | 0.89 ± 0.40 | -11.33 ± 15.54 |
|  | PsA SF +ADA | 1.76 ± 0.69 | 0.79 ± 0.12 | -4.97 ± 5.95 |
|  | OA SF | 1.18 ± 1.00 | 1.34 ± 0.19 | -0.19 ± 0.42 |
| **Psoriatic arthritis**<br>**explant tissues** | Baseline | 1.00 | 1.00 | 1.00 |
|  | PsA SF | 2.59 | 50.55 | 4.38 |

To evaluate whether OA explant tissues recapitulated transcriptional responses observed in PsA tissue, a single available PsA TKA donor with matched PsA SF was compared with OA tissue exposed to PsA SF. *COL1A1*, *COLX*, *ADAMTS5*, and *MMP1* in cartilage-bone tissue and *CD86*, *MMP1*, *CCL2*, and *IL1B* in synovium tissue showed similar patterns of expression between OA and PsA tissue treated with PsA SF (Fig. S4A-B). Together, these findings support the use of OA explant tissues exposed to PsA SF to recapitulate key transcriptional features observed in matched PsA tissue responses. PsA tissue treated with PsA SF also showed similar modest changes in IL-6 and pan MMP enzymatic activity; CCL2 changes were, however, more pronounced in PsA vs. OA tissue (Table 1).

### PsA SF induced different explant tissue behavior compared to OA SF

PsA SF treatment resulted in significant upregulation of *MMP3* compared to both untreated and OA SF-treated explant co-cultures (Fig. 2A). PCA of the osteochondral gene panel showed clear separation of OA explant co-cultures treated with PsA SF vs. OA SF and negative and positive control groups (Fig. 2B). Euclidean distance analysis with PsA SF, OA SF, or pro-inflammatory cytokines clustered 1.32, 1.48 and 6.46 units from the negative control, respectively.

**Figure 2.**
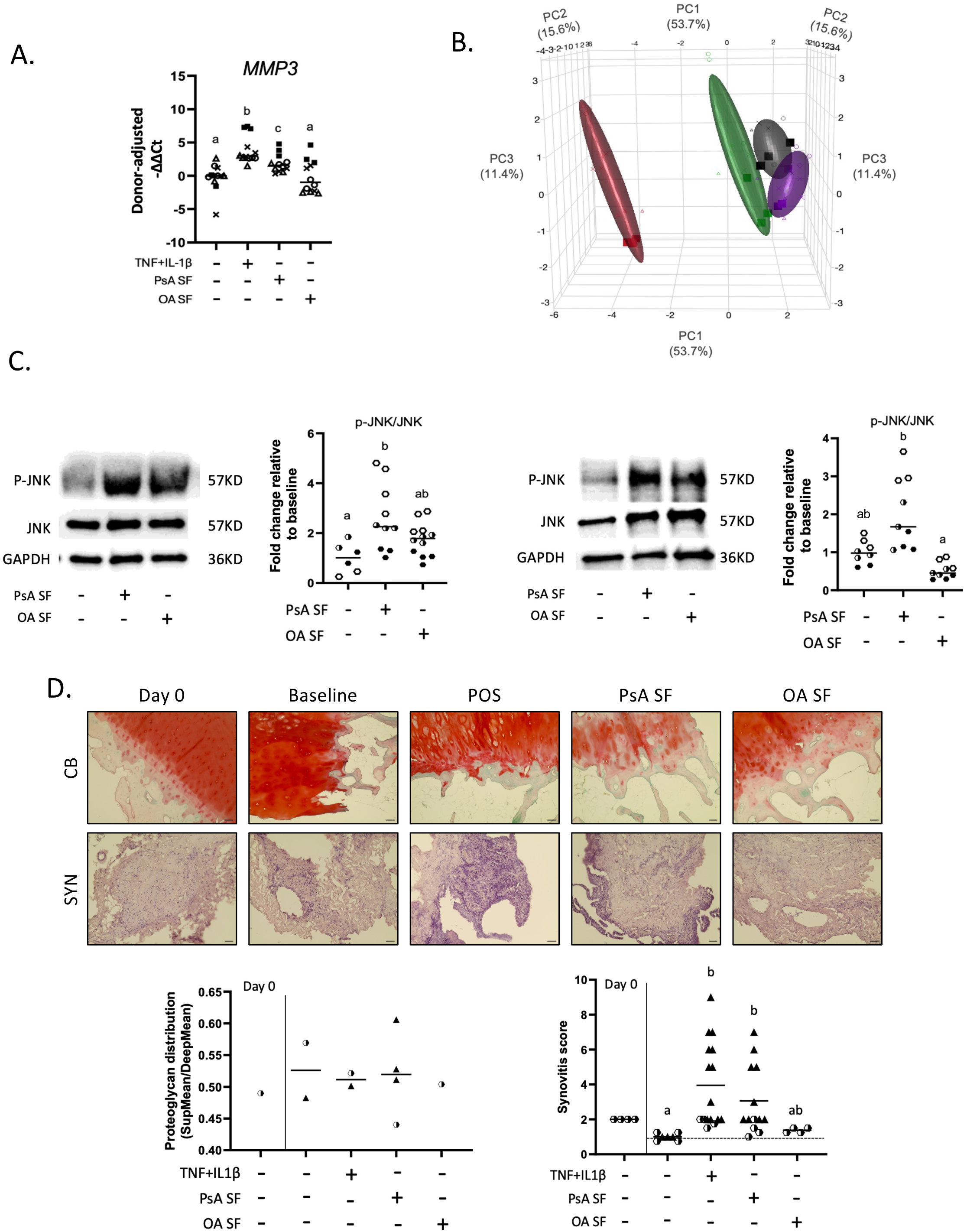
PsA SF induces different explant tissue behavior compared to OA SF. **(A)** Relative MMP3 expression (−ΔΔCt) across experimental conditions, normalized to baseline and adjusted for OA tissue donor effects (n = 4 donors, triplicate). **(B)** 3D PCA of CB gene expression profiles. Black, baseline; red, POS; green, PsA SF; purple, OA SF. PCA loading coefficients are provided (Suppl Table 6). **(C)** Representative western blots and densitometric quantification of p-JNK/JNK ratios in SYN (left) and CB (right) explants. Values were normalized to baseline. **(D)** Representative Safranin O/Fast Green-stained CB and H&E-stained SYN sections with corresponding quantification of proteoglycan distribution and synovitis scores. Synovitis scores were normalized to the matched baseline mean for each OA tissue donor (n = 2 OA tissue donors; n=4 PsA SF donors). Scale bars: 100 μm (CB), 50 μm (SYN). Symbols represent individual OA tissue donors (Suppl Table 1). Different letters indicate statistical significance (A and C-SYN: one-way ANOVA with Tukey’s post hoc test; C-CB and synovitis: Kruskal-Wallis test with Dunn’s multiple comparisons test; p < 0.05).

PC1 accounted for 53.7% of the total variance and showed clear separation between PsA SF and OA SF groups (shift = 2.24), capturing the primary axis of structural tissue integrity-related genes versus inflammatory and catabolic remodeling genes (Suppl Table 6). Signaling pathway analysis demonstrated significantly increased JNK activation in synovium explants following PsA SF treatment relative to baseline conditions and higher JNK activation in PsA vs. OA SF-treated explants (Fig. 2C).

Full-depth histological analysis did not reveal significant differences between cartilage proteoglycan distribution in superficial vs. deep zones across baseline (mean: 0.53), positive control (0.51), PsA (0.52) and OA SF treatment (0.50) (Fig. 2D). By contrast, synovium explants demonstrated significantly increased synovitis scores with positive and PsA SF treatment relative to baseline conditions (mean normalized Krenn scores: 4.18 and 3.06 vs 1.0, respectively; Fig. 2D). OA SF-treated explants demonstrated intermediate synovitis severity (1.38) that was not significantly different from baseline conditions.

After subtracting SF-alone effects and normalizing to untreated OA explant tissues, IL-6 and CCL2 showed modest increases across both PsA and OA SF conditions (Table 1). Pan-MMP activity, however, remained highly donor-variable and did not consistently align with transcriptional or histological changes (Table 1 & Suppl Table 4).

Overall, our model showed that PsA SF induced distinct osteochondral and synovium gene expression profiles, synovial inflammation, and JNK activation compared with OA SF.

### Explant tissue co-culture model treated with PsA SF demonstrated donor-specific sensitivity to DEX and ADA treatments

To test the utility of the model in screening drugs in a patient-responsive manner, we examined the PsA SF donor-driven variability in a subset of donors. Single OA donor-derived explant tissues were treated with PsA SF from three donors, to which either DEX, included as a broad anti-inflammatory control reflecting intra-articular glucocorticoid use in PsA (19), or the anti-TNF biologic ADA (20) was added (Fig. 3A).

**Figure 3.**
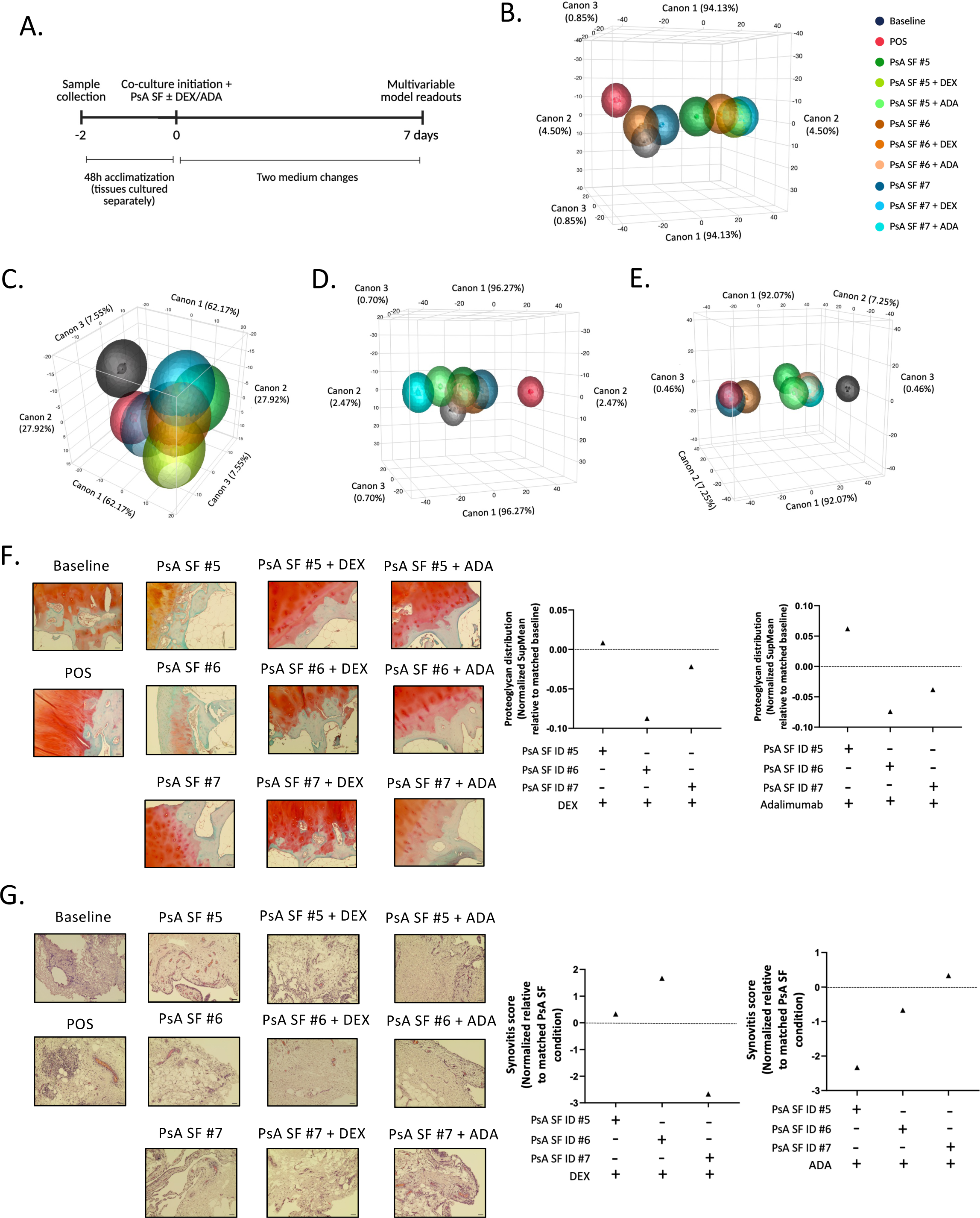
PsA SF donor-specific changes to DEX and ADA treatments in the explant model. **(A)** Experimental schematic of DEX or ADA treatment. **(B-E)** Canonical discriminant analysis of CB and SYN gene expression profiles following DEX **(B-C)** or ADA **(D-E)** treatment. Ellipses represent 50% probability contours. Darker colors indicate untreated PsA SF donor conditions, whereas lighter shades indicate matched DEX- or ADA-treated groups. Canonical structure coefficients are provided (Suppl Table 7). **(F-G)** Representative Safranin O/Fast Green-stained CB sections **(F)** and H&E-stained SYN sections **(G)** with corresponding changes in proteoglycan distribution and synovitis scores relative to matched PsA SF controls (n=1 OA tissue donor; n=3 PsA SF donors (#5-7); scale bars = 50 μm. Symbols correspond to individual PsA SF donors (Suppl Table 1).

Unsupervised cluster analysis showed PsA SF donor-driven gene expression profiles at baseline and after treatment with DEX (Fig 3B, C). Specifically, in osteochondral explanted tissues, PsA SF donor ID #6 demonstrated the largest DEX-induced shift (Euclidean distance = 35.39); PsA SF donor ID #7 had smaller DEX-induced shifts (26.04) followed by PsA SF donor ID #5 (17.71; Fig.3B). Canonical contribution analysis showed that treatment with DEX resulted in shifts along canonical axis 1 (94.13% variance) in osteochondral gene expression, primarily driven by *MMP1*, 19.14%; *RANKL*, 12.73%; *IL6*, 11.75%; *COLX*, 10.97%; *OPG*, 10.83%; *ADAMTS5*, 10.45% (Suppl Table 7).

Synovium tissue explants however showed that PsA SF donor ID #7 had the largest DEX-induced shift in gene expression (27.18), followed by PsA SF donor ID #5 (23.44); PsA SF donor ID #6 showed moderate changes (9.1; Fig.3C). Canonical axis 1 (62.17% variance) following DEX treatment was driven by inflammatory and macrophage-associated pathways, *MMP9*, 26.18%; *CD163*, 17.76%; *IL6*, 11.16%; *IL1B*, 10.68%; *IL23*, 9.38%.

Although different canonical gene contributors were observed following ADA treatment, with axis 1 dominated by *OPG* (17.68%), *MMP1* (15.68%), *IL6* (14.86%), *RANKL* (14.51%), and *MMP3* (13.98%) (Suppl. Table 7), PsA SF donor ID #6 remained the most sensitive (Euclidean distance = 17.56), followed by PsA SF donor ID #7 (8.71) and PsA SF donor ID #5 (7.74, Fig 3D). In synovium explants, ADA induced more modest responses, with Euclidean distances ranging from 8.54-9.92 across all donors (Fig. 3E). Canonical contribution analysis of synovium gene expression showed *IL1B*, 19.88%; *CCL2*, 17.79%; *MMP1*, 14.26%; *CXCL8*, 12.64% (Suppl Table 7) as key drivers. PsA SF donor ID #7 still remained most sensitive in synovium gene expression changes to ADA treatment; however, PsA SF donor IDs #6 was more sensitive than PsA SF donor ID #5 to ADA treatment; different from their respective gene expression changes to DEX treatments.

The model was thus sensitive enough to capture donor-specific differences in composite gene expression responses to DEX and ADA treatments. While osteochondral composite gene scores showed that all three PsA SF donors had similar changes to both DEX and ADA (PsA SF Donor ID# 6 > 7 > 5), synovium composite gene scores showed small differences between donors to DEX vs. ADA treatments (DEX: PsA SF Donor ID #7 > 5 > 6; ADA: PsA SF Donor ID #7 > 6 > 5). Thus, the model’s synovium tissue gene expression readouts captured differences between PsA SF donor changes to DEX and ADA.

The model also detected donor specific proteoglycan histological changes that were aligned with the donor-specific osteochondral gene changes. PsA SF Donor ID #6, which showed the greatest changes in osteochondral composite gene expression, exhibited the largest shifts in proteoglycan distribution with both DEX and ADA treatments. There was a redistribution towards a more balanced superficial:deep zone profile (−0.09 and −0.08, respectively; Fig 3F). Similarly, PsA SF Donor ID #7, which demonstrated intermediate osteochondral gene expression changes, showed modest changes in proteoglycan distribution, with DEX and ADA producing smaller reductions (−0.02 and −0.04, respectively; Fig 3F). PsA SF Donor ID #5, which showed the least change in osteochondral composite gene expression, exhibited minimal changes with DEX treatment (+0.01), while ADA resulted in a larger increase (+0.06), indicating some divergence between gene expression and matrix-level responses in this PsA SF donor.

Synovitis histology scores also varied by PsA SF donor (Fig 3G) but did not completely align with synovium composite gene scores. PsA SF Donor ID #7 showed the largest synovium gene expression changes to both DEX and ADA, and the greatest reduction in synovitis with DEX treatment (−2.67), but not with ADA (+0.33; Fig 3G). PsA SF Donor ID #6 showed the smallest synovium gene expression changes and worsening synovitis score changes to DEX treatment (+1.67), whereas ADA produced modest synovial gene changes together with modestly reduced synovitis scores (−0.23; Fig 3G). PsA SF Donor ID #5 showed modest synovium gene expression changes to DEX and ADA, but slightly increased synovitis with DEX (+0.33) and a strong reduction in synovitis with ADA (−2.34).

Protein analysis further confirmed PsA SF donor-dependent changes with DEX and ADA treatments (Suppl Table 4), although only CCL2 showed consistent reductions with ADA treatment; IL-6 and MMP activity demonstrated donor-to-donor variability.

### Model outputs correlate with baseline clinical features of PsA SF donors

To determine whether these model outputs reflected coordinated or independent aspects of PsA SF-induced pathology, Spearman correlations between the multivariable readouts were examined and showed limited correlations (Fig 4A). Proteoglycan distribution correlated inversely with synovitis scores and CCL2 levels, and with specific, but not composite osteochondral gene scores. Synovitis scores showed modest associations with CCL2 (ρ=0.83, p=0.04) and MMP activity (ρ=0.83, p=0.04) but not with synovium composite gene scores. Thus, our model captures multiple independent features of PsA SF-induced extracellular matrix (ECM), gene and protein changes in the explanted tissue.

**Figure 4.**
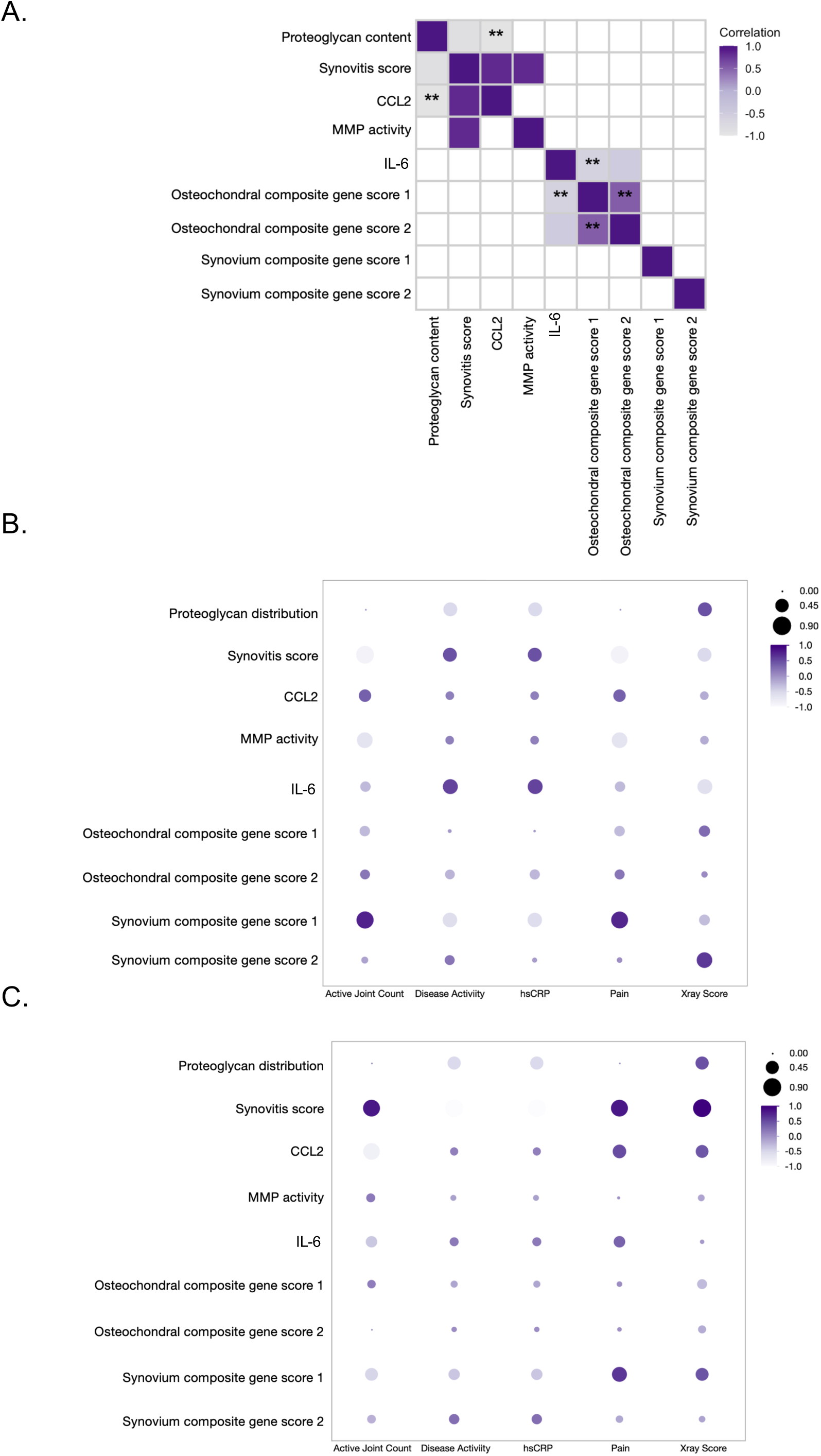
Model readouts correlate with baseline PsA donor clinical features. **(A)** Correlation matrix showing significant Spearman correlations (ρ) among explant model readouts under PsA SF treatment (n= 1 OA tissue donor; n = 3 PsA SF donors). Darker shading indicates stronger positive correlations; lighter shading indicates weaker or negative correlations. Double asterisks (**) indicate correlations that remained statistically significant following false discovery rate (FDR) correction. **(B-C)** Correlations between explant model readouts and baseline PsA SF donor clinical parameters following DEX **(B)** or ADA **(C)** treatment. Color indicates the Spearman correlation coefficient (ρ), with darker purple representing stronger positive correlations and lighter shading representing weaker or negative correlations. Dot size represents the absolute magnitude of the correlation coefficient (|ρ|). Clinical parameters included active joint count, PsA disease activity as measured by the Disease Activity Index for Psoriatic Arthritis (DAPSA), high-sensitivity C-reactive protein (hsCRP), pain, and X-ray score. Model-derived readouts in panels B and C represent treatment-induced changes relative to the matched PsA SF-treated condition, calculated as DEX-treated minus matched PsA SF-treated values (B) or ADA-treated minus matched PsA SF-treated values (C).

Model readouts in response to treatments were also examined in relation to PsA SF donor baseline clinical characteristics (Fig. 4B, C). For DEX-treated samples, synovium composite gene scores were significantly associated with active PsA-affected joint count (ρ=0.79, q=0.02) and baseline joint pain (ρ=0.76, q=0.02). Model synovitis scores following DEX treatment similarly showed strong correlations with baseline pain and active joint count (|ρ| ≥ 0.83), although these associations did not remain significant after multiple-testing correction.

In ADA-treated explant tissues, synovitis scores but not synovium composite scores were strongly associated with baseline clinical parameters, including disease activity, X-ray scores, and hsCRP (|ρ|≥ 0.83, q=0.001; Fig. 4C), demonstrating that histological synovial responses to ADA closely reflect clinical inflammatory burden.

PsA SF donor-specific treatment-associated changes were further compared with corresponding donor clinical histories of corticosteroid and TNF-targeting therapy response. Among all model readouts, histological synovitis scores showed the clearest qualitative agreement with clinical treatment history. PsA SF Donor ID#7, whose clinical history suggested initial improvements with steroid injections but no clear improvements with anti-TNF therapy, showed the largest reduction in model synovitis scores (4.67 to 2.0) to DEX but not ADA treatment. PsA SF Donor IDs #5 and #6 similarly, showed limited or inconsistent synovitis score changes with DEX and reduced scores with ADA treatments (4.67 to 2.33; 2.0 to 1.33, respectively), congruent with their respective clinical histories to corticosteroids or TNF-targeting therapies (Table 2 & Suppl Table 8).

**Table 2.** Explant model readouts and corresponding clinical notes for PsA SF donors. Model synovitis score changes following DEX or ADA treatment are summarized qualitatively as positive (+) or absent/negative (−), relative to PsA SF donor-treated conditions. Clinical notes provide qualitative summaries of treatment history and documented clinical observations for each PsA SF donor, based on available medical records. Model synovitis score readouts were based on N = 1 OA tissue donor exposed to N = 3 PsA SF donors, in duplicate or triplicate. PsA = psoriatic arthritis; SF = synovial fluid; DEX = dexamethasone; TNF = tumor necrosis factor; ADA = adalimumab; DMARD = disease-modifying antirheumatic drug

|  |  | **Donor 5** | **Donor 6** | **Donor 7** |
| --- | --- | --- | --- | --- |
| **Steroid Tx vs. Model changes to DEX** | **Clinical notes** | No documented steroid treatment or clinical benefit from steroid therapy | No clear clinical improvement following steroid injections | Initial clinical improvement following steroid injections, but benefit was not sustained |
|  | **Synovitis Score** | — | — | + |
| **Anti-TNF targeting Tx vs. Model changes to ADA** | **Clinical notes** | Clinical improvement observed with Otezla (PDE4 inhibitor), a cytokine-modulating therapy with indirect TNF suppression; direct anti-TNF clinical response unavailable | No documented Anti-TNF therapy or clinical benefit from Anti-TNF therapy | No clear clinical improvement following anti-TNF therapy; later improvement observed after switching to JAK inhibitor therapy |
|  | **Synovitis Score** | + | + | — |

Together, these findings support the capacity of model readouts (specifically synovitis scores) to recapitulate patient-specific changes to different anti-inflammatory treatments, supporting the potential utility of this model as a precision medicine platform.

## Discussion

We developed a PsA donor-specific, explanted osteochondral and synovium tissue-based end-stage joint model to evaluate multivariate responses to PsA SF over a 7-21-day period. The explant tissue-based transwell model integrated osteochondral and synovium tissue crosstalk, mediated by soluble factors. The model was sensitive to capture the PsA SF donor-driven heterogeneity indicated by the transcriptomic, histopathological, protein, and signaling pathway changes. Donor heterogeneity was also apparent when studying tissue changes in response to the corticosteroid dexamethasone or the anti-TNF drug adalimumab.

PsA SF induced distinct pro-inflammatory hallmark signatures, including increased *IL6, CCL2, CXCL8, MMP1, MMP3* and *WNT5A*, accompanied by altered cartilage proteoglycan distribution, moderate synovial activation, and increased JNK activation in the explanted tissues. Although only modest increases in IL-6 and CCL2 protein levels and no consistent secondary increases in pan-MMP activity beyond intrinsic SF levels were observed, secondary upregulation of pro-inflammatory and matrix-degrading genes in explanted joint tissues was evident. Thus, the model recapitulated intrinsic, primary PsA SF proteolytic effects together with secondary transcriptional responses, consistent with pathways associated with joint inflammation and damage in PsA. The multi-variable model readouts indicated selective coordination among specific features, as well as clear separation between distinct aspects of disease pathology. Synovitis scores associated with CCL2 and MMP activity and inversely with proteoglycan distribution scores, but not with synovium composite gene scores suggesting that the model captured a linked inflammatory-proteolytic axis and cartilage matrix integrity changes. The absence of model synovial composite gene correlations with other readouts is reflective of current gene panel curation; a broader panel would likely show stronger correlations and can be addressed in future model iterations.

Multivariable model readouts in response to drug treatments could be benchmarked against baseline PsA patient clinical indices including active (swollen or tender) joint counts, a measure of inflammatory burden. Changes in composite synovium gene expression, synovitis, and proteoglycan distribution in response to steroid treatment were associated with baseline active joint counts and patient-reported pain, indicating that the magnitude of synovial response reflected baseline disease activity. Given that ACR20/50/70 criteria, based on reduction in active joint counts, are often used to evaluate clinical responsiveness (21), these findings support the clinical relevance of our end-stage tissue model. By contrast, correlations between anti-TNF responses and baseline clinical indices, including hsCRP, were weaker and more variable, reflecting donor-dependent heterogeneity in response to TNF-targeting therapies. Baseline DAPSA (22–24) also correlated more consistently with steroid vs. anti-TNF treatment responses. The model was also useful in capturing PsA SF donor-specific changes in responses to two standard PsA treatments. Composite synovial gene expression, synovitis, and proteoglycan distribution scores showed concordant normalized responses to steroid treatment and showed preliminary alignment with PsA donor improvements to two anti-inflammatory treatments. Composite synovium gene scores, on the other hand, did not align with the histological synovitis scores with respect to changes induced by anti-TNF treatment. Notably, only synovitis scores aligned with PsA SF donor clinical improvements to TNF-targeting treatment. This suggests that synovitis scoring may represent a more sensitive and functionally relevant readout of anti-TNF induced changes within the current model. The current curated synovium gene panel, designed to capture key inflammatory and macrophage-associated pathways, may not fully reflect the broader cellular and signaling complexity of the synovial microenvironment (25). Expanding it to include additional SF cell populations and inflammatory components, such as Th17/Treg-associated cytokines, synovial fibroblast activation markers, and endothelial-related genes, may improve its ability to capture the anti-TNF-mediated effects and better reflect the multicellular components seen in PsA synovitis (23,26,27). Similarly, other model readouts, including chemokine levels and proteoglycan distribution, may more fully recapitulate donor-specific differential responses to corticosteroid or anti-TNF treatments. The variability in PsA SF donor-driven responses to corticosteroid and anti-TNF therapies mirrors clinical observations, where a proportion of PsA patients exhibit differential or non-responsiveness to these treatments (28–30).

Together, these findings support the ability of the model to capture donor-specific variability in therapeutic responses and highlight its potential to reflect clinically relevant treatment patterns within a controlled *ex vivo* system.

This is a first iteration of the model, and future versions could expand the synovium gene panel or incorporate cellular components into the PsA SF. Importantly, even with a small subset of PsA SF donors tested for responsiveness to both corticosteroid and anti-TNF treatments, this explant model was able to provide proof-of-concept data that it can (i) identify multivariable readouts associated with clinically relevant measures of disease burden and (ii) capture donor-specific treatment responses that aligned with patient clinical parameters. Thus, these proof-of-concept data support the potential of our patient-specific model for evaluating individualized responses to PsA therapies.

A limitation of our model is the use of primary joint tissues from OA rather than PsA patients. This approach leverages the convergence of key inflammatory signaling pathways, immune mediators, synovial inflammation, and cartilage remodeling processes observed in end-stage OA and severe PsA joint disease (23,31,32). Despite this, the model did demonstrate specificity to PsA SF donor-mediated changes and dictated changes in multivariate readouts in response to the drugs tested, mitigating concerns about using end-stage OA joint tissues. The limited availability of PsA patient tissue undergoing arthroplasty (14,33), relative to the substantially higher prevalence of osteoarthritis-related joint replacement, necessitated this approach. Notably, we showed that PsA joint tissue subjected to matched PsA SF had similar clustering based on selected osteochondral and synovial gene expression, although we acknowledge this was a single sample. Future model iterations could use induced pluripotent stem cell (iPSC)-derived joint cells (chondrocytes, synoviocytes) encapsulated in hydrogels (34–36); however, these stem cell derived models lack the complex ECM architecture that explant tissues possess (37).

The model used cell-free PsA SF, with variable cytokine, chemokine, and alarmin composition, likely drives distinct pathway activation in explanted tissues (38–40) but lacks immune effectors. This can be remedied by including complete PsA SF, which would necessitate fresh PsA SF to preserve cell viability. Alternatively, conditioned medium from PsA patient-derived Th17 or other immune cells (26,41) could be added to banked PsA SF to increase this dimensionality. Seven PsA SF donors were used to establish the model, and only a subset of three PsA SF donors were used to test PsA SF donor-driven differences to drug treatments. However, even within this small sample size, the model discerned donor differences that mimicked matched patient clinical parameters. We acknowledge the limited sample size and consider our model a proof of concept for further model development and validation.

In summary, our multi-compartmental, joint-specific explanted tissue model demonstrated PsA SF driven hallmarks of distinct inflammatory and degradative tissue changes; multivariable readouts of the model allowed exploration of different mechanisms of end-stage joint pathogenesis in response to cell-free PsA SF. Use of different PsA SF donors allowed the model to reflect PsA inter-donor variability and captured donor-specific differences to drug treatments and showed that model readouts were preliminarily aligned with the patient’s clinical changes. Taken together, this model can be leveraged to accelerate precision medicine approaches by capturing donor-specific variability in treatment responses.

## Acknowledgments

This work is supported by the Krembil Foundation Fund (UHN) Project Grant (No. 5790-7556-0713) to SV and VC. The work is also supported by the Schroeder Arthritis Institute via the University Health Network Foundation. VC is supported by the Dr. Dafna D. Gladman Chair in Psoriatic Arthritis Research, a joint Hospital-University Named Chair between the University of Toronto, the University Health Network, and the UHN Foundation. We thank the UHN Biobank, led by M. Kapoor and team, specifically K. Perry, C. Ward, and Luis Montoya from the Division of Orthopedics, for their work in OA sample management, consent, and data collection. We would also like to thank D. Pereira and D. Ganatra for their help with the collection and processing of the PsA patient samples. Schematics were developed using BioRender.com.

## Author contributions

A.Z. performed experiments, conducted all statistical analyses, generated figures, and drafted and revised the manuscript. M.R. performed experiments. K.O. conceived the study, contributed to manuscript editing and securing funding. V.C. secured funding and contributed to manuscript editing. S.V. conceived and supervised the study and contributed to writing and editing the manuscript. All authors reviewed and approved the final manuscript.

## References

1. FitzGerald O, Ogdie A, Chandran V, Coates LC, Kavanaugh A, Tillett W, et al. Psoriatic arthritis. Nat Rev Dis Primer. 2021 Aug 12;7(1):59. doi:10.1038/s41572-021-00293-y

2. Alinaghi F, Calov M, Kristensen LE, Gladman DD, Coates LC, Jullien D, et al. Prevalence of psoriatic arthritis in patients with psoriasis: A systematic review and meta-analysis of observational and clinical studies. J Am Acad Dermatol. 2019 Jan;80(1):251–265.e19. doi:10.1016/j.jaad.2018.06.027

3. Mease PJ, Gladman DD, Collier DH, Ritchlin CT, Helliwell PS, Liu L, et al. Etanercept and Methotrexate as Monotherapy or in Combination for Psoriatic Arthritis: Primary Results From a Randomized, Controlled Phase III Trial. Arthritis Rheumatol. 2019 Jul;71(7):1112–24. doi:10.1002/art.40851

4. Leung Y, Kavanaugh A, Ritchlin CT. Expert Perspective: Management of the Psoriatic Arthritis Patient After Failure of One Tumor Necrosis Factor Inhibitor. Arthritis Rheumatol. 2023 Aug;75(8):1312–24. doi:10.1002/art.42498

5. Baricza E, Marton N, Királyhidi P, Kovács OT, Kovácsné Székely I, Lajkó E, et al. Distinct In Vitro T-Helper 17 Differentiation Capacity of Peripheral Naive T Cells in Rheumatoid and Psoriatic Arthritis. Front Immunol. 2018 Apr 4;9:606. doi:10.3389/fimmu.2018.00606

6. Colucci S, Brunetti G, Cantatore F, Oranger A, Mori G, Quarta L, et al. Lymphocytes and synovial fluid fibroblasts support osteoclastogenesis through RANKL, TNFα, and IL-7 in an *in vitro* model derived from human psoriatic arthritis. J Pathol. 2007 May;212(1):47–55. doi:10.1002/path.2153

7. Lories RJ, Neerinckx B. Animal Models of Psoriasis and Psoriatic Arthritis. In: Adebajo A, Boehncke WH, Gladman DD, Mease PJ, editors. Psoriatic Arthritis and Psoriasis [Internet]. Cham: Springer International Publishing; 2016 [cited 2026 Jun 4]. p. 103–9. Available from: http://link.springer.com/10.1007/978-3-319-19530-8_12 doi:10.1007/978-3-319-19530-8_12

8. Sherlock JP, Joyce-Shaikh B, Turner SP, Chao CC, Sathe M, Grein J, et al. IL-23 induces spondyloarthropathy by acting on ROR-γt+ CD3+CD4−CD8− entheseal resident T cells. Nat Med. 2012 Jul;18(7):1069–76. doi:10.1038/nm.2817

9. Flores RR, Carbo L, Kim E, Van Meter M, De Padilla CML, Zhao J, et al. Adenoviral gene transfer of a single-chain IL-23 induces psoriatic arthritis–like symptoms in NOD mice. FASEB J. 2019 Aug;33(8):9505–15. doi:10.1096/fj.201900420R

10. Hagert C, Sareila O, Kelkka T, Jalkanen S, Holmdahl R. The Macrophage Mannose Receptor Regulate Mannan-Induced Psoriasis, Psoriatic Arthritis, and Rheumatoid Arthritis-Like Disease Models. Front Immunol. 2018 Feb 6;9:114. doi:10.3389/fimmu.2018.00114

11. Khmaladze I, Kelkka T, Guerard S, Wing K, Pizzolla A, Saxena A, et al. Mannan induces ROS-regulated, IL-17A–dependent psoriasis arthritis-like disease in mice. Proc Natl Acad Sci. 2014 Sep 2;111(35). doi:10.1073/pnas.1405798111

12. Robinson H, Kelly S, Pitzalis C. Basic Synovial Biology and Immunopathology in Psoriatic Arthritis. J Rheumatol Suppl. 2009 Aug 1;83(0):14–6. doi:10.3899/jrheum.090212

13. Chan MWY, Gomez-Aristizábal A, Mahomed N, Gandhi R, Viswanathan S. A tool for evaluating novel osteoarthritis therapies using multivariate analyses of human cartilage-synovium explant co-culture. Osteoarthritis Cartilage. 2022 Jan;30(1):147–59. doi:10.1016/j.joca.2021.09.007

14. Ciaffi J, Bianchi L, Di Martino A, Faldini C, Ursini F. Is Total Joint Arthroplasty an Effective and Safe Option for Psoriatic Arthritis Patients? A Scoping Review. J Clin Med. 2024 Sep 19;13(18):5552. doi:10.3390/jcm13185552

15. Oikonomopoulou K, Diamandis EP, Hollenberg MD, Chandran V. Proteinases and their receptors in inflammatory arthritis: an overview. Nat Rev Rheumatol. 2018 Mar;14(3):170–80. doi:10.1038/nrrheum.2018.17

16. Hui AY, McCarty WJ, Masuda K, Firestein GS, Sah RL. A systems biology approach to synovial joint lubrication in health, injury, and disease. WIREs Syst Biol Med. 2012 Jan;4(1):15–37. doi:10.1002/wsbm.157

17. Abji F, Rasti M, Gómez-Aristizábal A, Muytjens C, Saifeddine M, Mihara K, et al. Proteinase-Mediated Macrophage Signaling in Psoriatic Arthritis. Front Immunol. 2021 Mar 8;11:629726. doi:10.3389/fimmu.2020.629726

18. Krenn V, Morawietz L, Burmester G, Kinne RW, Mueller-Ladner U, Muller B, et al. Synovitis score: discrimination between chronic low-grade and high-grade synovitis. Histopathology. 2006 Oct;49(4):358–64. doi:10.1111/j.1365-2559.2006.02508.x

19. Rhen T, Cidlowski JA. Antiinflammatory Action of Glucocorticoids — New Mechanisms for Old Drugs. N Engl J Med. 2005 Oct 20;353(16):1711–23. doi:10.1056/NEJMra050541

20. Sundanum S, Orr C, Veale D. Targeted Therapies in Psoriatic Arthritis—An Update. Int J Mol Sci. 2023 Mar 28;24(7):6384. doi:10.3390/ijms24076384

21. Mease PJ, Gladman DD, Ritchlin CT, Ruderman EM, Steinfeld SD, Choy EHS, et al. Adalimumab for the treatment of patients with moderately to severely active psoriatic arthritis: Results of a double-blind, randomized, placebo-controlled trial. Arthritis Rheum. 2005 Oct;52(10):3279–89. doi:10.1002/art.21306

22. Schoels MM, Aletaha D, Alasti F, Smolen JS. Disease activity in psoriatic arthritis (PsA): defining remission and treatment success using the DAPSA score. Ann Rheum Dis. 2016 May;75(5):811–8. doi:10.1136/annrheumdis-2015-207507

23. Ritchlin CT, Colbert RA, Gladman DD. Psoriatic Arthritis. Longo DL, editor. N Engl J Med. 2017 Mar 9;376(10):957–70. doi:10.1056/NEJMra1505557

24. Coates LC, Helliwell PS. Treating to target in psoriatic arthritis: how to implement in clinical practice. Ann Rheum Dis. 2016 Apr;75(4):640–3. doi:10.1136/annrheumdis-2015-208617

25. Schonfeldova B, Zec K, Udalova IA. Synovial single-cell heterogeneity, zonation and interactions: a patchwork of effectors in arthritis. Rheumatology. 2022 Mar 2;61(3):913–25. doi:10.1093/rheumatology/keab721

26. Benham H, Norris P, Goodall J, Wechalekar MD, FitzGerald O, Szentpetery A, et al. Th17 and Th22 cells in psoriatic arthritis and psoriasis. Arthritis Res Ther. 2013 Sep 26;15(5):R136. doi:10.1186/ar4317

27. Celis R, Cuervo A, Ramírez J, Cañete JD. Psoriatic Synovitis: Singularity and Potential Clinical Implications. Front Med. 2019 Feb 11;6:14. doi:10.3389/fmed.2019.00014

28. Clunie G, McInnes IB, Barkham N, Marzo-Ortega H, Patel Y, Gough A, et al. Long-term effectiveness of tumour necrosis factor-α inhibitor treatment for psoriatic arthritis in the UK: a multicentre retrospective study. Rheumatol Adv Pract. 2018 Jul 1;2(2):rky042. doi:10.1093/rap/rky042

29. Keeling S, Maksymowych W. Difficult-To-Treat Psoriatic Arthritis Is Uncommon in a Real-World Registry. J Rheumatol. 2025 Jul;52(Suppl 2):114.1-114. doi:10.3899/jrheum.2025-0314.144

30. Neurath L, Sticherling M, Schett G, Fagni F. Targeting cytokines in psoriatic arthritis. Cytokine Growth Factor Rev. 2024 Aug;78:1–13. doi:10.1016/j.cytogfr.2024.06.001

31. Robinson WH, Lepus CM, Wang Q, Raghu H, Mao R, Lindstrom TM, et al. Low-grade inflammation as a key mediator of the pathogenesis of osteoarthritis. Nat Rev Rheumatol. 2016 Oct;12(10):580–92. doi:10.1038/nrrheum.2016.136

32. Scanzello CR, Goldring SR. The role of synovitis in osteoarthritis pathogenesis. Bone. 2012 Aug;51(2):249–57. doi:10.1016/j.bone.2012.02.012

33. Nystad TW, Husum YS, Furnes ON, Fevang BTS. Incidence and Predictive Factors for Orthopedic Surgery in Patients with Psoriatic Arthritis. J Rheumatol. 2018 Nov;45(11):1532–40. doi:10.3899/jrheum.180203

34. Nguyen D, Hägg DA, Forsman A, Ekholm J, Nimkingratana P, Brantsing C, et al. Cartilage Tissue Engineering by the 3D Bioprinting of iPS Cells in a Nanocellulose/Alginate Bioink. Sci Rep. 2017 Apr 6;7(1):658. doi:10.1038/s41598-017-00690-y

35. Huang J, Li A, Liang R, Wu X, Jia S, Chen J, et al. Future perspectives: advances in bone/cartilage organoid technology and clinical potential. Biomater Transl. 2024;5(4):425–43. doi:10.12336/biomatertransl.2024.04.007

36. Hu Y, Zhang H, Wang S, Cao L, Zhou F, Jing Y, et al. Bone/cartilage organoid on-chip: Construction strategy and application. Bioact Mater. 2023 Jul;25:29–41. doi:10.1016/j.bioactmat.2023.01.016

37. Banh L, Cheung KK, Chan MWY, Young EWK, Viswanathan S. Advances in organ-on-a-chip systems for modelling joint tissue and osteoarthritic diseases. Osteoarthritis Cartilage. 2022 Aug;30(8):1050–61. doi:10.1016/j.joca.2022.03.012

38. Gao W, McGarry T, Orr C, McCormick J, Veale DJ, Fearon U. Tofacitinib regulates synovial inflammation in psoriatic arthritis, inhibiting STAT activation and induction of negative feedback inhibitors. Ann Rheum Dis. 2016 Jan;75(1):311–5. doi:10.1136/annrheumdis-2014-207201

39. Kemble S, Croft AP. Critical Role of Synovial Tissue–Resident Macrophage and Fibroblast Subsets in the Persistence of Joint Inflammation. Front Immunol. 2021 Sep 3;12:715894. doi:10.3389/fimmu.2021.715894

40. Marsh L, Kemble S, Reis Nisa P, Singh R, Croft AP. Fibroblast pathology in inflammatory joint disease. Immunol Rev. 2021 Jul;302(1):163–83. doi:10.1111/imr.12986

41. He Y, Van Heeswijk B, Mus AMC, Davelaar N, Bisoendial R, Lubberts E. Active vitamin D acts in vitro as an adjuvant to anti-TNFα treatment in a psoriatic synovial fibroblast activation model by modulating human Th17 activity. RMD Open. 2025 Jul;11(3):e005547. doi:10.1136/rmdopen-2025-005547

42. Haltmayer E, Ribitsch I, Gabner S, Rosser J, Gueltekin S, Peham J, et al. Co-culture of osteochondral explants and synovial membrane as in vitro model for osteoarthritis. Ahmad R, editor. PLOS ONE. 2019 Apr 2;14(4):e0214709. doi:10.1371/journal.pone.0214709

43. Afara I, Singh S, Moody H, Oloyede A. A Comparison of the Histochemical and Image-Derived Proteoglycan Content of Articular Cartilage. Anat Physiol. 2013;03(02). doi:10.4172/2161-0940.1000120

44. Sun Y, Mauerhan DR, Kneisl JS, James Norton H, Zinchenko N, Ingram J, et al. Histological Examination of Collagen and Proteoglycan Changes in Osteoarthritic Menisci. Open Rheumatol J. 2012 Apr 19;6(1):24–32. doi:10.2174/1874312901206010024

45. Rasti M, Barazandeh AF, Robb KP, Low R, Kolade O, Fan K, et al. Mesenchymal Stromal Cells Immunosuppress Osteoarthritis Synovial Fluid Modulated Monocytes via IL-6 and CCL2 [Internet]. Immunology; 2025 [cited 2026 Jun 27]. Available from: http://biorxiv.org/lookup/doi/10.1101/2025.08.14.669875 doi:10.1101/2025.08.14.669875

