## Supplementary-File-1-Figures-and-Tables for "Joint tissue explant model using psoriatic arthritis synovial fluid as a tool to capture patient-specific responses to treatments"

**Supplementary Table 1. Characteristics of OA explant tissue donors and PsA synovial fluid donors.**

Plot symbols shown for OA tissue donors correspond to those used in principal component analysis (PCA), discriminant analysis (DA), and other donor-level graphical data presentations. Alphabetical donor IDs correspond to those used in protein data tables. Figure numbers preceded by "S" indicate supplementary figures. Clinical parameters, including active and swollen joint counts (66/68 method), Disease Activity using DAPSA score, patient-reported pain, and X-ray scores, were obtained from patient records under REB approval (#21-5591). Peripheral blood lymphocyte, monocyte, neutrophil, and WBC counts are reported as $\times10^9$/L. Reference ranges were as follows: lymphocytes, 1.5-4 $\times10^9$/L; monocytes, 0.2-0.8 $\times10^9$/L; neutrophils, 2-7.5 $\times10^9$/L; and WBCs, 4-11 $\times10^9$/L. OA = osteoarthritis; PsA = psoriatic arthritis; SF = synovial fluid; TKA = total knee arthroplasty; BMI = body mass index; DAPSA = Disease Activity in Psoriatic Arthritis; hsCRP = high-sensitivity C-reactive protein; WBC = white blood cell; M = male; F = female; NA = not available.

**Supplementary Table 2. Characteristics of OA SF donors.**

Peripheral blood lymphocyte, monocyte, neutrophil, and WBC counts are reported as $\times10^9$/L. Reference ranges were as follows: lymphocytes, 1.5-4 $\times10^9$/L; monocytes, 0.2-0.8 $\times10^9$/L; neutrophils, 2-7.5 $\times10^9$/L; and WBCs, 4-11 $\times10^9$/L. OA = osteoarthritis; SF = synovial fluid; M = male; F = female; BMI = body mass index; WBC = white blood cell; NA = not available.

**Supplementary Table 3. Curated Cartilage-bone and synovium gene panels and primers for qPCR.**

Forward and reverse primer sequences are shown in the 5'→3' direction. CB = cartilage-bone; SYN = synovium.

**Supplementary Table 4. IL-6, CCL2, and MMP activity levels in conditioned medium**

Values are presented as raw donor-level values following double normalization to matched SF-alone conditions and baseline controls. Donor letter annotations correspond to OA tissue donor IDs (Suppl Table 1).

**Supplementary Table 5. PC loading coefficients for cartilage-bone and synovium gene expression.**

Bold values indicate strong contributors (|loading coefficient| ≥ 0.7). PCA loading coefficients represent the direction and relative contribution of each gene to the PCs. Relative contribution (%) for each gene within a PC was calculated as the squared loading coefficient divided by the sum of squared loading coefficients for all genes, multiplied by 100. The percent variance explained by each PC is indicated in Fig. 1C.

**Supplementary Table 6. PC loading coefficients for cartilage-bone gene expression.**

Bold values indicate strong contributors (|loading coefficient| ≥ 0.7). PCA loading coefficients represent the direction and relative contribution of each gene to the PCs. Relative contribution (%) for each gene within a PC was calculated as the squared loading coefficient divided by the sum of squared loading coefficients for all genes, multiplied by 100. The percent variance explained by each principal component is indicated in Fig. 2B.

**Supplementary Table 7. Canonical structure coefficients for discriminant analysis across treatments in cartilage-bone and synovium compartments.**
Bold values indicate strong contributors to canonical axes ($|r| \geq 0.7$). Canonical structure coefficients represent the correlation between each gene and the canonical axes. Relative canonical contribution (%) was calculated as the squared structure coefficient ($r^2$) divided by the sum of squared structure coefficients for all genes, multiplied by 100. The percent variance explained by each canonical axis is indicated in Fig. 3B-E.

**Supplementary Table 8. Comparison of donor-specific model readouts and clinical treatment notes for PsA SF donors following corticosteroid and anti-TNF treatments.**
Model readout changes following DEX or ADA treatment are summarized qualitatively as positive (+), minimal (~), or absent/negative (-) relative to untreated PsA SF donor conditions. Clinical notes summarize documented treatment history and clinical observations for each PsA SF donor.

**Supplementary Figure 1. Original Western blot images for JNK signaling proteins in synovium and cartilage-bone explants.**
**(A)** Uncropped Western blot images of phosphorylated JNK (p-JNK), total JNK, and GAPDH in synovium (SYN) explants cultured under baseline, PsA synovial fluid (PsA SF), and OA synovial fluid (OA SF) conditions. **(B)** Uncropped Western blot images of phosphorylated JNK (p-JNK), total JNK, and GAPDH in cartilage-bone (CB) explants cultured under baseline, PsA SF, and OA SF conditions. GAPDH served as the loading control.

**Supplementary Figure 2. Timepoint effects on transcriptional profiles in the co-culture model**
**(A-B)** PCA of osteochondral (CB; **A**) and synovial (SYN; **B**) gene expression profiles across co-culture timepoints following exposure to PsA SF. Each point represents one donor-derived co-culture sample. Black, COCUL-BASE; red, COCUL-POS; green, COCUL-PsA. Distinct symbols indicate culture timepoints: day 7 (circle), day 14 (triangle), and day 21 (square). Ellipses represent group clustering distributions. The percent variance explained by each PC axis is indicated. The right panels show PCA loading vectors representing gene contributions to the PC separation. Genes projecting in similar directions are positively associated, whereas genes projecting in opposite directions contribute inversely to sample separation. OA tissue donor samples were analyzed in triplicate (Suppl Table 1).

**Supplementary Figure 3. PsA synovial fluid dose-dependent effects on transcriptional profiles in the co-culture model.**
**(A-B)** PCA of CB **(A)** and SYN **(B)** gene expression profiles across PsA SF concentrations. Each point represents one donor-derived co-culture sample. Black, baseline; red, POS; green, 30% PsA SF; orange, 40% PsA SF; and blue, 50% PsA SF. Ellipses represent group clustering distributions. The percent variance explained by each PC axis is indicated. The right panels show PCA loading vectors representing gene contributions to the PC separation. Distinct symbols correspond to individual OA tissue donors (N = 3 OA tissue donors, in triplicate), as defined in Suppl Table 1.

**Supplementary Figure 4. Comparison of selected gene expression responses between OA tissue and PsA tissue exposed to PsA synovial fluid.**

**(A-B)** Scatter plots comparing relative gene expression (-ΔΔCt) of selected genes between OA tissue + PsA SF and PsA tissue + PsA SF conditions in **(A)** CB and **(B)** synovium. Each point represents the mean -ΔΔCt value of an individual gene under the indicated condition. OA tissue + PsA SF values represent the mean -ΔΔCt across OA donor-derived co-culture samples (n=3 OA tissues; n=3 PsA SF donors), whereas PsA tissue + PsA SF values represent the mean -ΔΔCt from technical replicates derived from a single matched PsA tissue and PsA SF donor. Red diagonal lines indicate theoretical equivalence between OA tissue + PsA SF and PsA tissue + PsA SF responses.

### Supplementary Table 1.

| Information about the Osteoarthritis tissues | | | | | | | | | | | | | | | | |
|---|---|---|---|---|---|---|---|---|---|---|---|---|---|---|---|---|
| Donor ID | - | - | A | B | - | - | - | - | C | - | D | E | - | - | - | F |
| Plot symbols | Y | ● | △ | ■ | - | - | ○ | □ | X | ★ | ◆ | ◇ | ⬣ | ⬡ | [illegible] | ▲ |
| Used in figure | S2, S3 | 1, S3 | 1, 4A, S2 | 1, 2A-B, 4A | S2 | S2 | 4A, S3 | 4A, S2 | 1, 2A-B, 4A, S 2 | 1, 4A | 1, 4A | 1, 4A | 2C, S1 | 2C, S1 | 2C, S1 | 2D, 3B-G, 4 |
| Age | 66 | 66 | 66 | 58 | 84** | 48 | 59 | 69 | 72 | 65 | 86 | 70 | 63 | 61 | 77 | 74 |
| Sex | M | F | M | M | M** | M | F | M | F | M | F | M | M | F | F | F |
| BMI (Kg/m$^2$) | 29.3 | 25.4 | 28.1 | 37.1 | 24.4** | 26.7 | 29.8 | NA | 23 | 26.6 | 21.6 | 39.5 | 26.6 | 33.7 | 30.8 | 33 |
| Lymphocyte | 1.8 | 2.5 | 1.3 | 1.3 | 1.9** | 2.1 | 2.9 | 1.7 | 2.3 | 1.4 | 1.3 | 2.1 | 1.2 | 1.9 | 3.1 | 2.1 |
| Monocyte | 0.6 | 0.6 | 0.6 | 0.6 | 2.5** | 0.5 | 0.6 | 0.7 | 0.6 | 0.4 | 0.7 | 0.8 | 0.6 | 0.8 | 0.7 | 0.8 |
| Neutrophil | 5.7 | 4.3 | 5.6 | 4.2 | 8.6** | 4 | 3.9 | 5.2 | 4.7 | 2.9 | 4.4 | 4.4 | 4.2 | 3.3 | 5.3 | 5 |
| WBC | 8.3 | 7.5 | 7.6 | 6.4 | 13** | 6.8 | 7.6 | 7.8 | 7.9 | 4.8 | 6.5 | 7.6 | 6.2 | 6.4 | 9.3 | 8.4 |

| Information about the Psoriatic arthritis synovial fluids and tissue* | | | | | | | |
|---|---|---|---|---|---|---|---|
| PsA SF ID | 1 | 2 | 3 | 4* | 5 | 6 | 7 |
| Age | 28 | 39 | 56 | 64 | 55 | 62 | 66 |
| BMI | 28.86 | 21.16 | 24.16 | 22.5 | 25.31 | 31.1 | 26.99 |
| Number of active and swollen joints | 6 | 1 | 2 | 2 | 1 | 1 | 2 |
| DAPSA score | 22 | 10 | 16 | 13.6 | 26.01 | 35.3 | 14.92 |
| Patient pain | 1 | 3 | 9 | 4 | 3 | 3 | 7 |
| X-ray score | 0 | 0 | 2 | 0 | 10 | 0 | 14 |
| hsCRP (mg/L) | 7.32 | 4.34 | 1.46 | 2.6 | 17.01 | 27.3 | 0.92 |

*PsA SF donor ID#4 provided matched PsA tissue and synovial fluid.
** Clinical information for this donor was obtained following total knee arthroplasty.

#### Supplementary Table 2.

| OA SF ID | Age | Sex | BMI | Lymphocyte | Monocyte | Neutrophil | WBC |
|---|---|---|---|---|---|---|---|
| 8 | NA | NA | NA | NA | NA | NA | NA |
| 9 | 63 | M | 25.93 | 1.5 | 0.5 | 4.8 | 7.1 |
| 10 | 73 | M | NA | 1.4 | 0.7 | 4.5 | 6.9 |
| 11 | 70 | F | 28.94 | 2.2 | 0.7 | 4.0 | 7.0 |
| 12 | 63 | M | 41.51 | 2.9 | 0.5 | 4.3 | 7.9 |
| 13 | 76 | M | 29.36 | 1.3 | 0.4 | 2.7 | 4.5 |
| 14 | 83 | M | 32.24 | 1.2 | 0.4 | 1.8 | 3.6 |

### Supplementary Table 3

| Gene | Protein Alias | Panel | Forward | Reverse | Reported Functionality |
|---|---|---|---|---|---|
| ***RPL13A*** | Ribosomal protein L13a | CB | TGCGACAAAACCTCCTCCTT | TGTTGATGCCTTCACAGCGTA | Reference |
| ***TBP*** | TATA-binding box protein | CB | GAGACGAGTTCCAGCGCAA | GCAGCAAACCGCTTGGGAT | Reference |
| ***ACAN*** | Aggrecan | CB | ACTCTGGGTTTTCGTGACTCT | ACACTCAGCGAGTTGTCATGG | Anabolic matrix |
| ***ADAMTS5*** | ADAM metallopeptidase with thrombospondin type 1 motif 5 (aggrecanase 2) | CB | TGGCTCACGAAATCGGACATT | TGCATTTGGACCAGGGCTTA | Catabolic matrix |
| ***BGLAP*** | Bone Gamma-Carboxyglutamate Protein (osteocalcin) | CB | CTCACACTCCTCGCCCTATTG | GCTTGGACACAAAGGCTGCAC | Osteoblast–Osteoclast Signaling Regulators |
| ***CCL2*** | C-C motif chemokine ligand 2 (monocyte chemotactic factor) | CB | TCGCCTCCAGCATGAAAGTC | GGTGACTGGGGCATTGATTG | Chemotaxis & adhesion |
| ***COL1A1*** | Collagen I | CB | CCTGGATGCCATCAAAGTCT | CGCCATACTCGAACTGGAAT | Anabolic matrix |
| ***COL2A1*** | Collagen II | CB | CATGAGGGCGCGGTAGAG | CCAGCCTCCTGGACATCCT | Anabolic matrix |
| ***COLX*** | Collagen X | CB | TGCTGCCACAAATACCCTTT | GTGGACCAGGAGTACCTTGC | Anabolic matrix |
| ***FGF18*** | Fibroblast growth factor 18 | CB | GTACGTGGGCTTCACCAAGA | TGGTCACCGTCGTGTACTTG | Anabolic matrix |
| ***IL6*** | Interleukin 6 | CB | GTCCAGTTGCCTTCTCCCTG | ACCAGGCAAGTCTCCTCATTG | Inflammation |
| ***MMP3*** | Matrix metallopeptidase 3 (stromelysin 1) | CB | CAAAGCTTCAGTGTTGGCTG | GGCCAGGGATTAATGGAGAT | Catabolic matrix |
| ***OPG*** | Osteoprotegrin | CB | GCGCTCGTGTTTCTGGACA | AGTATAGACACTCGTCACTGGTG | Osteoblast–Osteoclast Signaling Regulators |
| ***PRG4*** | Proteoglycan 4 (lubricin) | CB | AGGCCCCATGTGTTCATGC | GCGCAAAGTAGTCAGTCCATCT | Anabolic matrix |
| ***RANKL/TNFSF11*** | TNF Superfamily Member 11 | CB | GCGTCGCCCTGTTCTTCTAT | TGCAGTGAGTGCCATCTTCTG | Osteoblast–Osteoclast Signaling Regulators |
| ***SOX9*** | SRY-box transcription factor 9 | CB | ATGCAAGCATGTGTCATCCA | AGGTCTGTCAGTGGGCTGAT | Anabolic matrix |
| ***SPP1*** | Secreted phosphoprotein 1 (osteopontin) | CB | GCCGAGGTGATAGTGTGGTT | TGAGGTGATGTCCTCGTCTG | Anabolic matrix |
| ***WNT5a*** | WNT family member 5A | CB | GCCAGTATCAATTCCGACATCG | TCACCGCGTATGTGAAGGC | Osteoblast–Osteoclast Signaling Regulators |
| ***B2M*** | Beta-2-microgloblin | CB, SYN | GAGGCTATCCAGCGTACTCCAAAG | GCTGCTTACATGTCTCGATCCCAC | Reference |
| ***MMP1*** | Matrix metallopeptidase 1 (collagenase 1) | CB, SYN | CTCAATTTCACTTCTGTTTTCTG | CATCTCTGTCGGCAAATTCGT | Catabolic matrix |
| ***ACTB*** | Actin Beta | SYN | CTCACCATGGATGATGATATCGC | AGGAATCCTTCTGACCCATGC | Reference |
| ***GAPDH*** | Glyceraldehyde-3-Phosphate Dehydrogenase | SYN | ACAACTTTGGTATCGTGGAAGG | GCCATCACGCCACAGTTTC | Reference |
| ***CD163*** | CD163 Molecule | SYN | TTTGTCAACTTGAGTCCCTTCAC | TCCCGCTACACTTGTTTTCAC | Macrophage polarization |
| ***CD206 (MRC1)*** | CD206 Molecule | SYN | CTACAAGGGATCGGGTTTATGGA | TTGGCATTGCCTAGTAGCGTA | Macrophage polarization |
| ***CD86*** | CD86 Molecule | SYN | CCATCAGCTTGTCTGTTTCATTCC | GCTGTAATCCAAGGAATGTGGTC | Macrophage polarization |
| ***IL17A*** | Interleukin 17A | SYN | AGATTACTACAACCGATCCACCT | GGGGACAGAGTTCATGTGGTA | Inflammation |
| ***IL1B*** | Interleukin 1 beta | SYN | GTACCTGTCCTGCGTGTTGA | GGGAACTGGGCAGACTCAAA | Inflammation |
| ***IL23A*** | Interleukin 23 Subunit Alpha | SYN | GACAACAGTCAGTTCTGCTTGC | AGAGAAGGCTCCCCTGTGAA | Inflammation |
| ***IL8/CXCL8*** | C-X-C Motif Chemokine Ligand 8 | SYN | AAATTTGGGGTGGAAAGGTT | TCCTGATTTCTGCAGCTCTGT | Chemotaxis & adhesion |
| ***MMP9*** | Matrix Metallopeptidase 9 | SYN | AGACCTGGGCAGATTCCAAAC | CGGCAAGTCTTCCGAGTAGT | Catabolic matrix |

#### Supplementary Table 4.

| Condition | OA Donor IDs | IL-6 | CCL2 | MMP activity |
|---|---|---|---|---|
| | | Raw (ng/mL) | Raw (ng/mL) | Raw (RFU) |
| Baseline | D | 1.00 | 1.00 | 1.00 |
| | E | 1.00 | 1.00 | 1.00 |
| | C | 1.00 | 1.00 | 1.00 |
| | A | 1.00 | 1.00 | 1.00 |
| | B | 1.00 | 1.00 | 1.00 |
| | F | 1.00 | 1.00 | 1.00 |
| Pro-inflammatory cytokines | D | 12.85 | 9.65 | 5.43 |
| | E | 12.72 | 21.00 | 3.86 |
| | C | 33.91 | 9.01 | 3.60 |
| | A | 17.68 | 9.96 | 8.23 |
| | B | 18.58 | 12.45 | 79.07 |
| | F | 30.62 | 6.26 | 31.75 |
| PsA SF | D | 1.46 | 1.45 | 0.78 |
| | E | 1.06 | 1.65 | 0.83 |
| | C | 2.30 | 1.11 | -0.95 |
| | A | 1.40 | 1.75 | -2.77 |
| | B | 1.78 | 1.52 | -26.70 |
| | F (+PsA SF #5) | 2.82 | 1.06 | 2.30 |
| | F (+PsA SF #6) | 2.32 | 2.00 | -10.60 |
| | F (+PsA SF #7) | 2.19 | 1.84 | -10.65 |
| PsA SF + DEX | C | 0.92 | 0.88 | -1.42 |
| | A | 1.23 | 1.44 | -2.88 |
| | B | 0.95 | 1.31 | -41.51 |
| | F (+PsA SF #5) | 1.98 | 0.56 | -0.05 |
| | F (+PsA SF #6) | 1.34 | 0.63 | -11.09 |
| | F (+PsA SF #7) | 2.01 | 0.51 | -11.00 |
| PsA SF + ADA | D | 1.36 | 0.94 | -1.44 |
| | E | 0.95 | 0.71 | -0.20 |
| | F (+PsA SF #5) | 2.75 | 0.81 | -0.33 |
| | F (+PsA SF #6) | 1.68 | 0.63 | -12.01 |
| | F (+PsA SF #7) | 2.07 | 0.84 | -10.90 |
| OA SF | C | 2.33 | 1.19 | 0.06 |
| | A | 0.66 | 1.55 | 0.05 |
| | B | 0.54 | 1.29 | -0.67 |

**Supplementary Table 5.**

| Cartilage-Bone | | | | | | |
|---|---|---|---|---|---|---|
| Gene | PC1 loading coefficient | PC1 contribution (%) | PC2 loading coefficient | PC2 contribution (%) | PC3 loading coefficient | PC3 contribution (%) |
| *BGLAP* | **0.95** | 12.74 | 0.20 | 1.45 | -0.05 | 0.13 |
| *ACAN* | **0.93** | 12.24 | 0.17 | 1.06 | 0.09 | 0.51 |
| *IL6* | **-0.90** | 11.61 | 0.07 | 0.18 | 0.26 | 4.19 |
| *COL2A1* | **0.83** | 9.80 | -0.25 | 2.35 | 0.13 | 1.04 |
| *COL1A1* | **0.81** | 9.28 | 0.23 | 1.90 | 0.01 | 0.00 |
| *PRG4* | **0.78** | 8.59 | 0.03 | 0.03 | 0.33 | 6.78 |
| *SPP1* | **0.77** | 8.38 | 0.39 | 5.50 | 0.13 | 1.02 |
| *MMP1* | -0.65 | 5.95 | 0.56 | 11.74 | 0.18 | 2.15 |
| *COLX* | 0.64 | 5.85 | -0.37 | 4.92 | 0.45 | 12.92 |
| *MMP3* | -0.61 | 5.32 | 0.34 | 4.17 | 0.44 | 12.23 |
| *WNT5A* | -0.56 | 4.48 | 0.67 | 16.66 | 0.22 | 2.97 |
| *RANKL* | 0.18 | 0.48 | 0.65 | 15.70 | 0.48 | 14.81 |
| *ADAMTS5* | -0.20 | 0.56 | -0.63 | 14.63 | 0.56 | 20.21 |
| *SOX9* | 0.57 | 4.69 | 0.61 | 13.81 | -0.02 | 0.03 |
| *OPG* | -0.03 | 0.01 | -0.40 | 5.91 | 0.58 | 21.02 |
| Synovium | | | | | | |
| Gene | PC1 loading coefficient | PC1 contribution (%) | PC2 loading coefficient | PC2 contribution (%) | PC3 loading coefficient | PC3 contribution (%) |
| *CD206* | **0.93** | 17.06 | -0.11 | 0.58 | 0.13 | 1.79 |
| *CXCL8* | **0.88** | 15.16 | 0.19 | 1.86 | -0.31 | 10.69 |
| *MMP1* | **0.86** | 14.63 | 0.24 | 2.80 | 0.14 | 2.21 |
| *IL1B* | **0.85** | 14.22 | 0.36 | 6.50 | -0.08 | 0.65 |
| *CCL2* | **0.77** | 11.58 | -0.27 | 3.78 | -0.15 | 2.71 |
| *IL23* | **0.74** | 10.73 | 0.42 | 8.84 | -0.23 | 6.07 |
| *CD86* | 0.67 | 8.80 | -0.54 | 14.60 | 0.35 | 13.99 |
| *CD163* | 0.35 | 2.48 | -0.84 | 35.55 | 0.21 | 5.02 |
| *IL17A* | 0.34 | 2.29 | 0.51 | 13.33 | 0.62 | 43.53 |
| *MMP9* | -0.39 | 3.06 | 0.49 | 12.17 | 0.34 | 13.32 |

**Supplementary Table 6.**

| Cartilage-Bone | | | | | | |
|---|---|---|---|---|---|---|
| Gene | PC1 loading coefficient | PC1 contribution (%) | PC2 loading coefficient | PC2 contribution (%) | PC3 loading coefficient | PC3 contribution (%) |
| *IL6* | **-0.97** | 11.59 | 0.17 | 1.17 | -0.03 | 0.06 |
| *ACAN* | **0.92** | 10.41 | 0.26 | 2.82 | 0.07 | 0.31 |
| *BGLAP* | **0.91** | 10.34 | 0.10 | 0.40 | 0.07 | 0.29 |
| *COL1A1* | **0.89** | 9.92 | 0.00 | 0.00 | -0.23 | 3.01 |
| *WNT5a* | **-0.89** | 9.88 | 0.27 | 3.06 | 0.20 | 2.28 |
| *PRG4* | **0.81** | 8.21 | 0.31 | 3.99 | 0.06 | 0.25 |
| *COL2A1* | **0.80** | 8.04 | 0.33 | 4.75 | 0.07 | 0.27 |
| *SOX9* | **0.71** | 6.24 | 0.45 | 8.59 | -0.38 | 8.50 |
| *SPP1* | **0.71** | 6.19 | 0.09 | 0.34 | 0.13 | 0.92 |
| *MMP1* | **-0.70** | 6.15 | 0.43 | 7.93 | -0.09 | 0.52 |
| *OPG* | 0.13 | 0.22 | **0.75** | 23.90 | -0.43 | 10.84 |
| *MMP3* | -0.57 | 4.05 | 0.59 | 15.11 | -0.47 | 13.05 |
| *ADAMTS5* | -0.49 | 3.02 | 0.57 | 13.98 | 0.24 | 3.43 |
| *COLX* | 0.56 | 3.89 | 0.33 | 4.55 | **0.70** | 29.00 |
| *RANKL* | -0.39 | 1.85 | 0.47 | 9.40 | 0.68 | 27.29 |

**Supplementary Table 7.**

| | Cartilage-Bone | | | | | | | |
|---|---|---|---|---|---|---|---|---|
| | DEX | | | | ADA | | | |
| Gene | Canon1 structure coefficients | Canon1 (%) | Canon2 structure coefficients | Canon2 (%) | Canon1 structure coefficients | Canon1 (%) | Canon2 structure coefficients | Canon2 (%) |
| *COL1A1* | 0.05 | 0.05 | -0.10 | 0.40 | -0.20 | 0.75 | -0.64 | 9.79 |
| *COL2A1* | -0.21 | 0.97 | 0.33 | 3.95 | 0.03 | 0.01 | **0.73** | 12.91 |
| *COLX* | **-0.72** | 10.97 | 0.30 | 3.39 | -0.02 | 0.01 | 0.49 | 5.71 |
| *MMP3* | -0.38 | 3.01 | **-0.78** | 22.24 | **0.85** | 13.98 | 0.15 | 0.54 |
| *MMP1* | **-0.95** | 19.14 | -0.16 | 0.94 | **0.90** | 15.68 | 0.26 | 1.58 |
| *PRG4* | -0.18 | 0.66 | -0.05 | 0.09 | 0.27 | 1.39 | 0.02 | 0.01 |
| *SPP1* | -0.14 | 0.43 | 0.26 | 2.50 | 0.05 | 0.05 | 0.47 | 5.38 |
| *ADAMTS5* | **-0.70** | 10.45 | 0.29 | 3.09 | 0.26 | 1.28 | **0.88** | 18.98 |
| *RANKL* | **-0.77** | 12.73 | -0.37 | 5.05 | **0.87** | 14.51 | 0.33 | 2.65 |
| *BGLAP* | -0.56 | 6.57 | 0.62 | 13.90 | -0.37 | 2.68 | 0.52 | 6.61 |
| *SOX9* | -0.34 | 2.47 | -0.40 | 5.80 | 0.64 | 7.80 | 0.42 | 4.25 |
| *IL6* | **-0.74** | 11.75 | -0.60 | 13.35 | **0.88** | 14.86 | -0.20 | 1.01 |
| *ACAN* | 0.31 | 2.10 | 0.45 | 7.51 | -0.14 | 0.40 | **0.80** | 15.57 |
| *FGF18* | -0.47 | 4.73 | -0.32 | 3.87 | 0.58 | 6.57 | 0.37 | 3.27 |
| *OPG* | **-0.71** | 10.83 | -0.62 | 13.92 | **0.96** | 17.68 | -0.03 | 0.02 |
| *WNT5a* | -0.38 | 3.14 | 0.00 | 0.00 | 0.35 | 2.34 | **0.70** | 11.73 |
| | Synovium | | | | | | | |
| | DEX | | | | ADA | | | |
| *Gene* | Canon1 structure coefficients | Canon1 (%) | Canon2 structure coefficients | Canon2 (%) | Canon1 structure coefficients | Canon1 (%) | Canon2 structure coefficients | Canon2 (%) |
| *MMP9* | **-0.86** | 26.18 | -0.27 | 2.88 | 0.25 | 1.51 | **-0.80** | 51.34 |
| *MMP1* | -0.49 | 8.33 | 0.47 | 8.76 | **-0.77** | 14.26 | 0.13 | 1.34 |
| *CD206* | -0.14 | 0.71 | **0.70** | 19.68 | -0.65 | 10.31 | -0.01 | 0.02 |
| *CD163* | **0.71** | 17.76 | 0.14 | 0.82 | -0.08 | 0.16 | 0.59 | 28.00 |
| *CD86* | -0.25 | 2.22 | 0.52 | 10.71 | -0.46 | 5.16 | 0.02 | 0.02 |
| *IL1B* | -0.55 | 10.68 | 0.69 | 19.02 | **-0.91** | 19.88 | -0.23 | 4.28 |
| *IL17A* | 0.19 | 1.23 | 0.19 | 1.43 | -0.25 | 1.52 | 0.33 | 8.52 |
| *IL23* | -0.52 | 9.38 | 0.44 | 7.55 | -0.54 | 7.06 | -0.23 | 4.36 |
| *CCL2* | -0.38 | 5.05 | 0.59 | 13.87 | **-0.86** | 17.79 | 0.04 | 0.11 |
| *CXCL8* | -0.46 | 7.30 | 0.39 | 6.05 | **-0.72** | 12.64 | -0.15 | 1.85 |
| *IL6* | -0.56 | 11.16 | 0.48 | 9.24 | -0.63 | 9.71 | -0.05 | 0.17 |

#### Supplementary Table 8.

| Treatment | Synovial fluid Donor | Model readouts | Change in model readouts | Clinical notes |
|---|---|---|---|---|
| Corticosteroid vs. Model changes to DEX | Donor 5 | Synovitis score | – | No documented steroid treatment or clinical benefit from steroid therapy |
| | | Synovium Composite gene score (PC1) | + | |
| | | Osteochondral Composite gene score (PC1) | + | |
| | | Proteoglycan content | ~ | |
| | | Protein (CCL2 dominant) | + | |
| | Donor 6 | Synovitis score | – | No clear clinical improvement following steroid injections |
| | | Synovium Composite gene score (PC1) | – | |
| | | Osteochondral Composite gene score (PC1) | + | |
| | | Proteoglycan content | + | |
| | | Protein (CCL2 dominant) | + | |
| | Donor 7 | Synovitis score | + | Initial clinical improvement following steroid injections, but benefit was not sustained |
| | | Synovium Composite gene score (PC1) | + | |
| | | Osteochondral Composite gene score (PC1) | + | |
| | | Proteoglycan content | ~ | |
| | | Protein (CCL2 dominant) | + | |
| Adalimumab | Donor 5 | Synovitis score | + | Clinical improvement observed with Otezla (PDE4 inhibitor), a cytokine-modulating therapy with indirect TNF suppression; direct anti-TNF clinical response unavailable |
| | | Synovium Composite gene score (PC1) | ~ | |
| | | Osteochondral Composite gene score (PC1) | + | |
| | | Proteoglycan content | + | |
| | | Protein (CCL2 dominant) | + | |
| | Donor 6 | Synovitis score | + | No documented Anti-TNF therapy or clinical benefit from Anti-TNF therapy |
| | | Synovium Composite gene score (PC1) | + | |
| | | Osteochondral Composite gene score (PC1) | + | |
| | | Proteoglycan content | + | |
| | | Protein (CCL2 dominant) | + | |
| | Donor 7 | Synovitis score | – | No clear clinical improvement following anti-TNF therapy; later improvement observed after switching to JAK inhibitor therapy |
| | | Synovium Composite gene score (PC1) | + | |
| | | Osteochondral Composite gene score (PC1) | + | |
| | | Proteoglycan content | – | |
| | | Protein (CCL2 dominant) | + | |

**Supplementary figure 1.**

A.

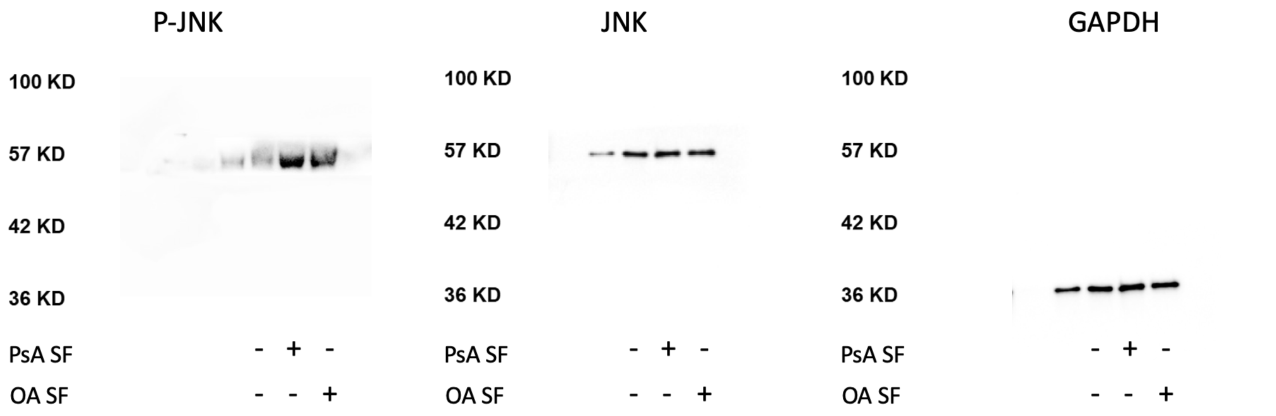


B.

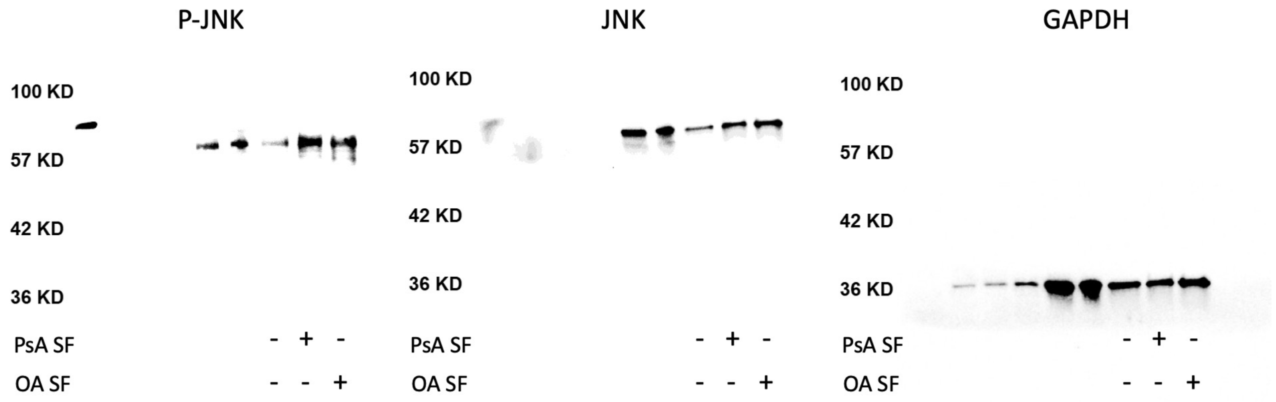

**Supplementary figure 2.**

A.

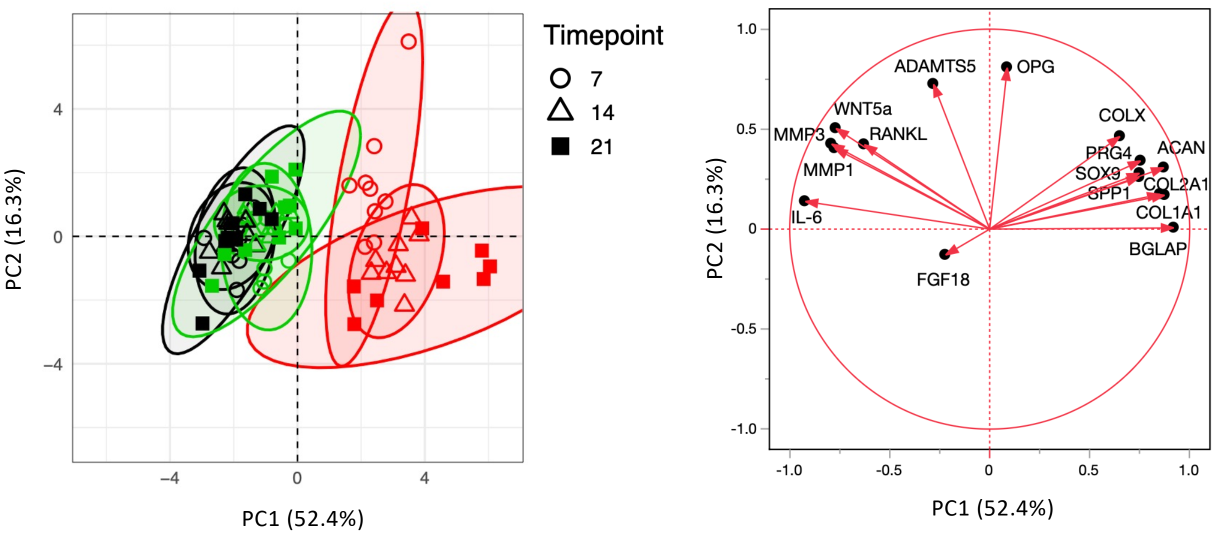


B.

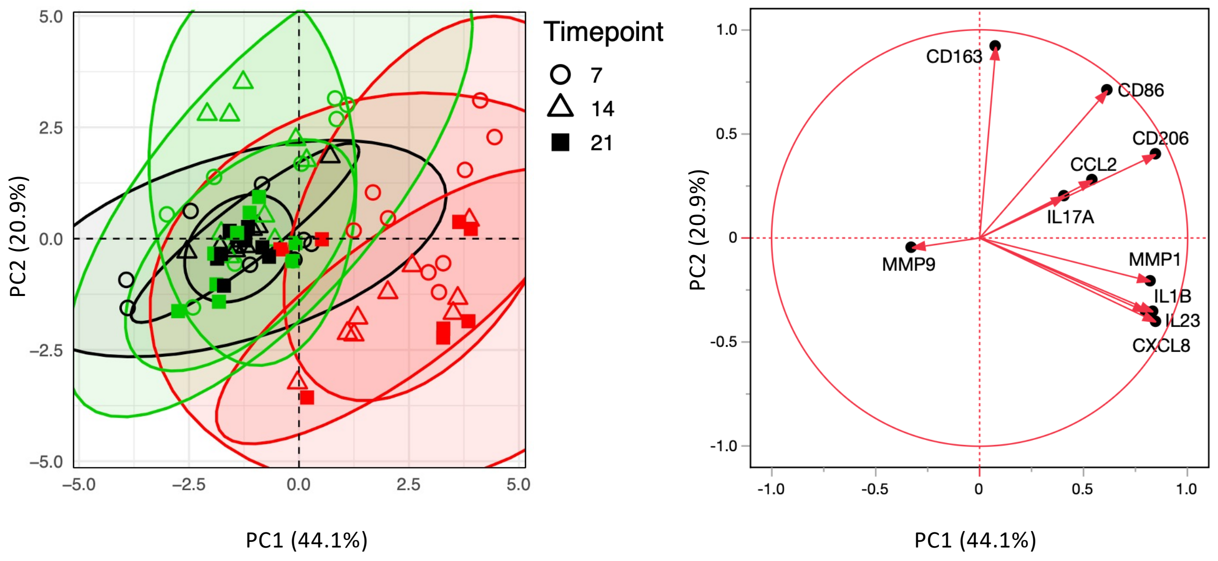

**Supplementary figure 3.**

A.

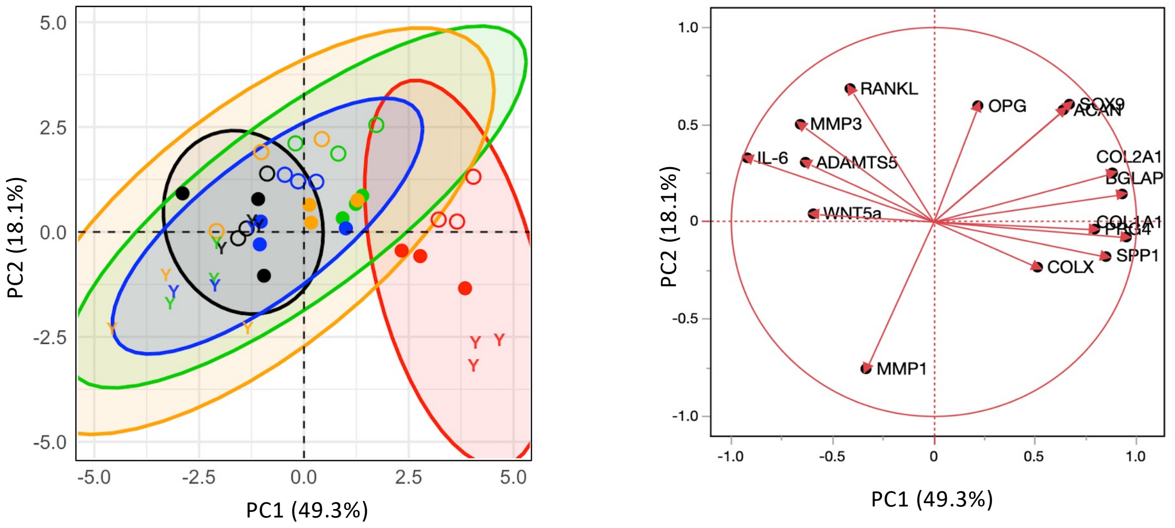


B.

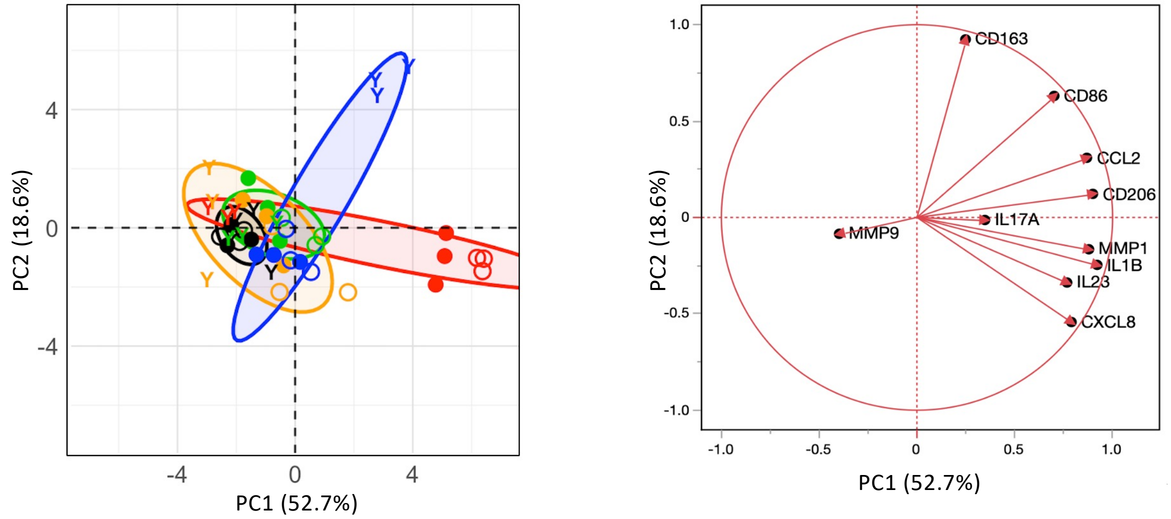

**Supplementary figure 4.**

A.

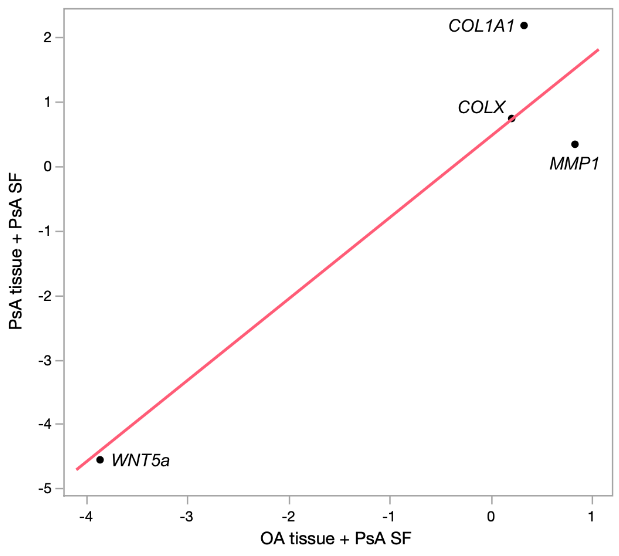


B.

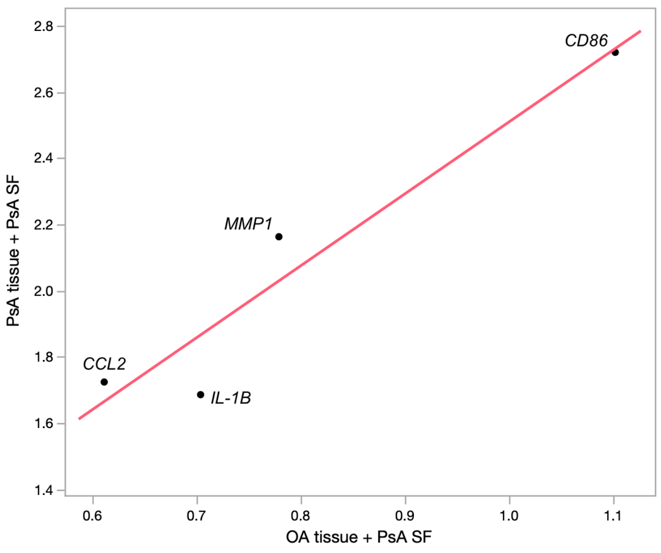
