## Supplementary-File-2-Supplementary-Methods for "Joint tissue explant model using psoriatic arthritis synovial fluid as a tool to capture patient-specific responses to treatments"

### Detailed Explant Processing and Co-culture Conditions

Cartilage-bone explants were generated using a 2.7 mm osteochondral grafting system (Smith & Nephew) as previously described (42) and placed in 24-well plates. Synovium tissues were trimmed, minced, and cultured on 0.4 µm transwell inserts at a 2:1 cartilage-bone: synovium ratio (13). Explants were maintained in optimized co-culture medium following a 48-hour acclimatization period.

Experimental conditions included: medium control, TNF (10 ng/mL) + IL-1β (10 ng/mL) positive control, 30% OA SF (Osteoarthritis synovial fluid), 30% PsA SF (Psoriatic arthritis synovial fluid), PsA SF + dexamethasone (DEX, 100 nM), and PsA SF + adalimumab (ADA, 2.5 µg/mL). Explants were cultured for up to 21 days with media changes performed twice weekly.

### Histological Quantification

Safranin O/Fast Green staining intensity was quantified in FIJI (ImageJ software, National Institutes of Health, Bethesda, MD, USA), using standardized regions of interest (ROIs) placed in superficial and deep cartilage zones (43–44) (Suppl Fig M1). Three ROIs were placed within each cartilage zone and averaged to obtain representative superficial and deep staining intensities for each section. Normalized superficial proteoglycan distribution was then calculated as Superficial / (Superficial + Deep) to assess relative proteoglycan localization across cartilage zones.

Synovium sections stained with H&E were scored using the Krenn synovitis scoring system, generating total synovitis scores ranging from 0-9 (18).

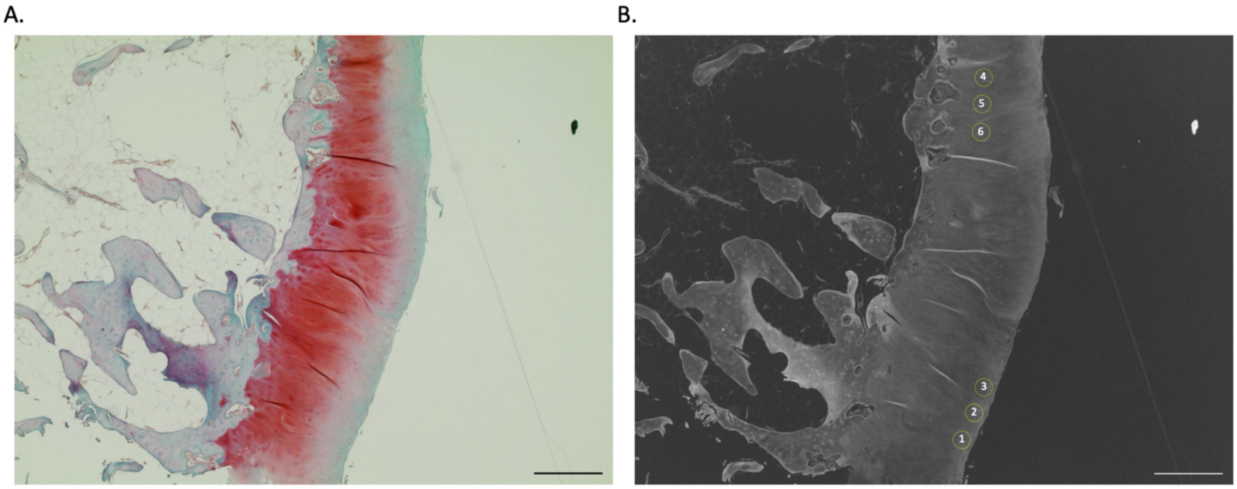


**Supplementary Figure M1. ROI placement for quantification of cartilage proteoglycan distribution.** (A) Representative Safranin O/Fast Green-stained cartilage-Bone section. (B) ROI placement used for quantification of superficial (ROIs 1–3) and deep (ROIs 4–6) cartilage zones. Three ROIs per zone were averaged and used to calculate normalized superficial proteoglycan distribution as Superficial / (Superficial + Deep). Circular ROIs diameter = 63.8 µm, Scale bars = 500 µm

**Cytokine and MMP Quantification**

IL-6 and CCL2 concentrations in conditioned medium were quantified using ELISA MAX Deluxe Set (BioLegend) and Human CCL2/MCP-1 ELISA Kit (Bio-Techne), respectively. Total MMP activity was measured using the SensoLyte® 520 Generic MMP Activity Kit (Anaspec) according to the manufacturer's instructions. The medium from triplicate wells was pooled prior to analysis.

**Western Blot Analysis**

Western blotting was performed as previously described (45) to assess total c-Jun N-terminal kinases (JNK; Cell Signaling Technology) and phosphorylated JNK (ABclonal). Band intensities were quantified and normalized to GAPDH.

**Statistical and Multivariate Analysis**

Normality was assessed using the Shapiro-Wilk test. Normally distributed gene expression datasets were analyzed using one-way ANOVA followed by Tukey's honestly significant difference (HSD) post hoc testing with OA tissue donor included as a blocking factor. Non-parametric datasets were analyzed using the Kruskal-Wallis test, followed by Steel-Dwass or Dunn's post hoc multiple comparisons tests, as appropriate. Paired comparisons within matched PsA SF donor conditions were assessed using Wilcoxon signed-rank tests.

Gene expression values were normalized to respective baseline co-culture controls and housekeeping genes. Soluble mediator levels were corrected for intrinsic cytokine and proteolytic activity by subtracting matched SF-alone values at equivalent dilutions prior to normalization against untreated baseline co-culture controls.

Associations between model-derived readouts and donor clinical parameters were assessed using Spearman's rank correlation ($\rho$). Due to limited sample size, covariate adjustment was not performed.

Principal component analysis (PCA) loading coefficients were used to assess the contributions of variables to the principal components. Canonical discriminant analysis (DA) structure coefficients ($r$) were used to evaluate variable correlations with canonical axes. Canonical contribution (%) was calculated by dividing each squared structure coefficient ($r^2$) by the total sum of squared coefficients within the corresponding canonical axis. Euclidean distances between group centroids were calculated from principal component scores using summed squared differences across components.
